# A niche-dependent redox rheostat regulates epithelial stem cell fate

**DOI:** 10.1101/2025.01.26.634856

**Authors:** Xi Chen, Krishnan Raghunathan, Bin Bao, Elsy Ngwa, Andrew Kwong, Zhongyang Wu, Stephen Babcock, Clara Baek, George Ye, Anoohya Muppirala, Qianni Peng, Michael Rutlin, Mantu Bhaumik, Daping Yang, Daniel Kotlarz, Unmesh Jadhav, Meenakshi Rao, Eranthie Weerapana, Xu Zhou, Jose Ordovas-Montanes, Scott B. Snapper, Jay R. Thiagarajah

## Abstract

The niche environment surrounding intestinal stem cells (ISCs) varies along the length of intestine and provides key cues that regulate stem cell fate. Here, we investigated the role of cellular redox balance in colonic ISC function. We show that hypoxia and Wnt signaling synergize to restrict the reactive oxygen species (ROS) generating enzyme NADPH oxidase 1 (NOX1) to the crypt base in the distal colon. NOX1 function maintains a more oxidative cell state that licenses cell cycle entry, altering the balance of asymmetric stem cell self-renewal and directing lineage commitment. Mechanistically, cell redox state directs a self-reinforcing circuit that connects hypoxia inducible factor 1 (HIF1α)-dependent signaling with regulation of the metabolic enzyme isocitrate dehydrogenase 1 (IDH1). Our studies show that cellular redox balance is a central and niche-dependent regulator of epithelial homeostasis and regeneration and provide a basis for understanding disease propensity in the distal large intestine.

**GRAPHICAL ABSTRACT:** 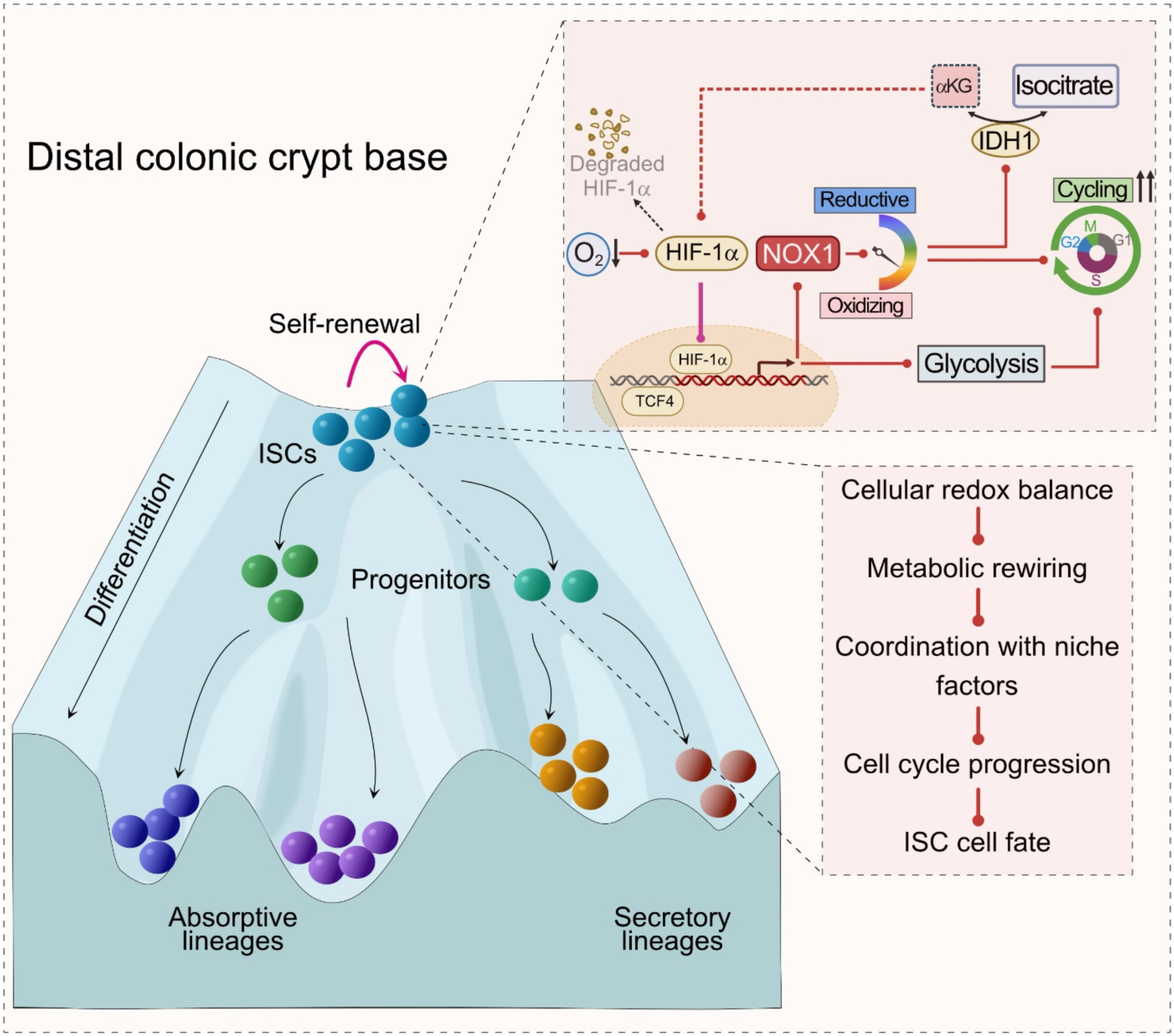

**HIGHLIGHTS:**

- The balance of cycling intestinal stem cells (ISCs) versus committed epithelial cells in the uniquely hypoxic niche of the distal colon is regulated by NADPH oxidase 1 (NOX1) dependent H_2_O_2_ both at homeostasis and during regeneration.
- Physiological increase in cellular H_2_O_2_ favors maintenance of glycolysis in ISCs for self-renewal through regulation of isocitrate dehydrogenase 1 activity.
- Maintenance of the increased cellular oxidative state stabilizes HIF1α through a re-enforcing metabolic circuit.
- A shift from a relatively oxidative to a reductive cell environment in distal colonic ISCs leads to decreased progression through the cell cycle and altered cell fate determination.

## INTRODUCTION

The maintenance, renewal and response to stress of post-natal tissues relies on the activity of compartment-specific multi-potent stem cells.^1,2^ The intestinal epithelium undergoes rapid constant renewal that is dynamically regulated by stem cells that reside in the base of crypt glands. In both the small and large intestine, cycling stem cells, marked by high expression of the leucine-rich repeat-containing G-protein coupled receptor 5 (Lgr5), control epithelial cell density and the generation of various differentiated epithelial cell lineages.^3^ Disruption of the balance between self-renewal and differentiation results in a wide range of diseases from developmental disorders to cancer.

Stem cell replication for either self-renewal or progenitor cell population generation is dependent on efficient progression through the cell cycle and requires environment-specific metabolic adaptations for energy and biomass production. Metabolic changes in intestinal epithelial stem-cells (ISCs) are well appreciated to have a critical role in regulating proliferation and differentiation.^4^ The niche environment surrounding all tissue resident stem cells, including ISCs, provides key variables that shape metabolic state, including nutrient provision, oxygen availability and developmental signals such as Wnt.^5^ Most studies to-date have focused on ISC metabolism in the context of maintenance and regeneration in the small intestine.^6–10^ The intestine however exhibits considerable differences in niche environment along its cephalo-caudal axis. The mammalian distal colon is a relatively unique mucosal environment with major differences in blood supply, oxygen tension and nutrient use from the small intestine as well as from the proximal colon.^11,12^ In humans the distal colon also represents a relatively fragile environment as suggested by the observation that major disease states such as inflammatory bowel disease or cancer are concentrated or in some case originate at this tissue site.

Several lines of evidence suggest that ISC metabolism is characterized by relative reliance on aerobic glycolysis and a lower fraction of oxidative phosphorylation with important contributions from neighboring secretory epithelial cells.^7,13,14^ An important factor in maintenance and switching of metabolic state in proliferating cells in other tissues is cellular redox balance, with oxidative metabolism reliant on the generation of both reductive and oxidative equivalents in the form of NADH or NAD+ respectively. Redox signaling and balance in relation to stem cell function has been linked to metabolic homeostasis in other tissues including pluripotent stem cells.^15,16^ However, how cytosolic redox balance in the context of regional niche-specific cues such as oxygen availability, affects intestinal stem cell metabolic status and functions in homeostatic intestinal epithelial renewal and regeneration remains poorly understood.

Here we investigated the role of a major colonic epithelial reactive oxygen species (ROS) producing enzyme, NADPH oxidase 1 (NOX1) in tissue maintenance and regeneration. We show that the unique spatial restriction of NOX1 expression to the distal versus proximal colon crypt base results in a compartment specific role in regulating stem cell renewal, and the balance between proliferation and differentiation in both homeostasis and active regeneration. Using cell-specific redox and stem cell reporter mice, along with single-cell transcriptomics, cysteine proteomics, and cell metabolic studies we show that NOX1 maintains cytosolic redox balance to enable glycolytic metabolism, efficient cell cycle progression and favor distal colonic ISC self-renewal versus differentiation.

## RESULTS

### NOX1 controls cytosolic redox status in crypt base stem cells in the distal colon

Several NADPH oxidase enzymes have been shown to be functionally important in intestinal epithelial cells along the gastrointestinal tract including NOX1, NOX4 and DUOX2.^17,18^ Previous studies have shown increasing *Nox1* expression in colonic versus small intestine mucosal tissue^19^ and have suggested increased expression within colonic stem cells.^20^ We therefore sought to precisely spatially characterize *Nox1* expression in WT mice. We found that *Nox1* is very specifically expressed in distal versus proximal colonic crypt base epithelial cells (Figures 1A-C and S1A). Within distal colonic crypt base cells, *Nox1* expression co-localized with *Lgr5^+^* cells, in contrast to DUOX2 which is largely expressed in colonic surface epithelial cells (Figures 1D and S1B). To quantitatively assess the intracellular redox status in intestinal epithelial cells (IECs), we constructed a cell-specific redox reporter mouse (Rosa26-roGFP2-Orp1^flox/flox^) (Figures S1C and S1D), where the fluorescence ratio (405 nm and 488 nm) measures relative cytosolic hydrogen peroxide (H_2_O_2_) concentration^21^ and crossed this with Villin-Cre to generate an IEC-specific hydrogen peroxide reporter mouse (Figure S1E). To validate the probe dynamic range and assess levels of oxidized versus reduced probe, colon tissues were treated with either the oxidant diamide or reducing agent dithiothreitol (DTT) (Figures S1F and S1G). Consistent with the spatial restriction of *Nox1* the crypt base in the distal colon is relatively more oxidized than the upper crypt as well as the crypt tip (Figures 1E and 1F). Comparison of WT;roGFP2-Orp1^Vil^ to NOX1^KO^;roGFP2-Orp1^Vil^ showed significantly lower cytosolic H_2_O_2_ and relative less cellular oxidation in distal colonic crypt base cells when NOX1 is deficient (Figures 1G and 1H). Colonoids generated from the distal colon of NOX1^KO^ redox-reporter mice similarly showed significantly lower steady-state relative cellular H_2_O_2_ (Figures 1I-L). These data along with the observation that the expression of other NOX enzymes was unchanged in NOX1^KO^ mice (Figure S1H) suggest that NOX1 controls cytosolic redox status in crypt base stem cells.

**Figure 1.**
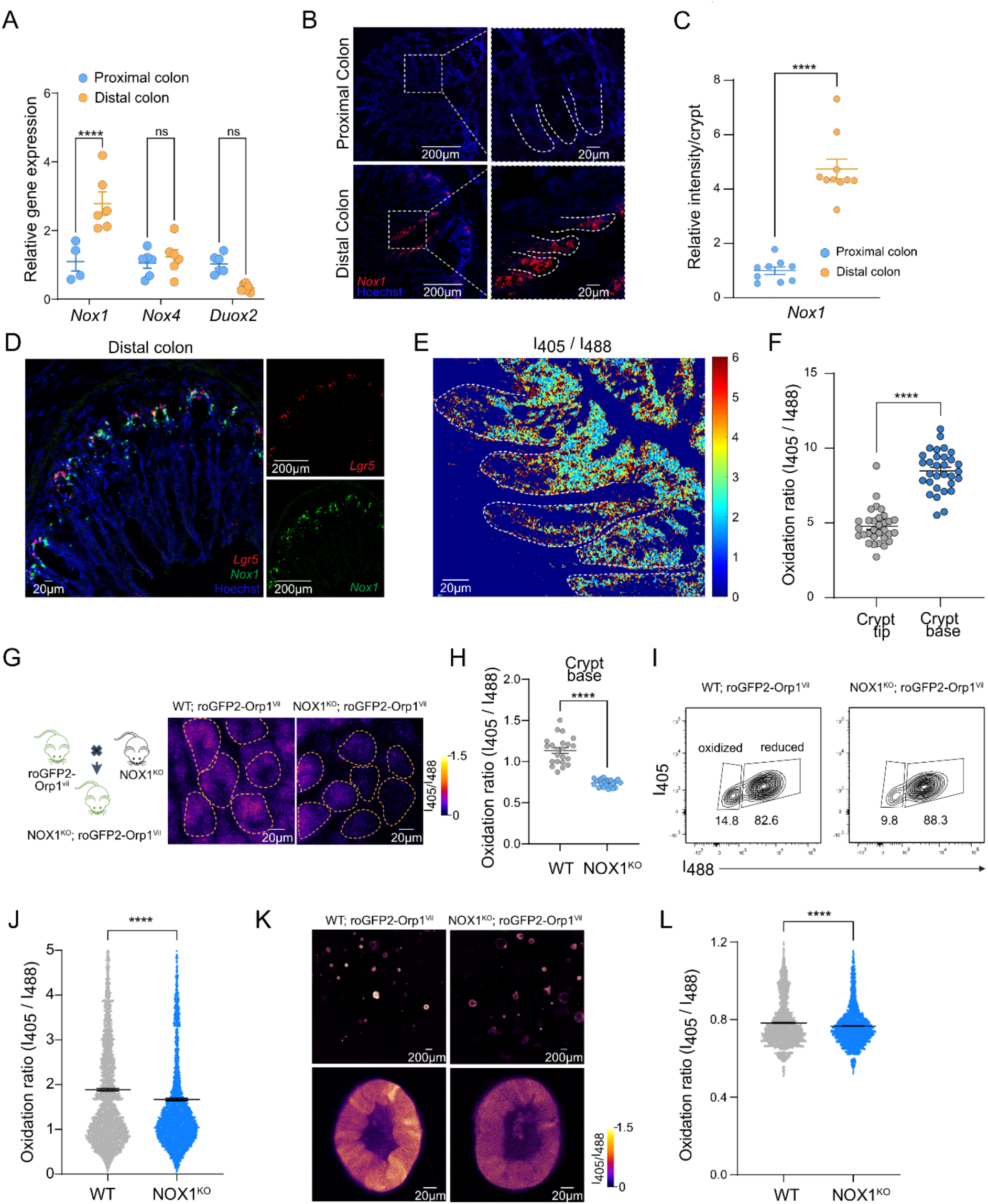
NOX1 controls cytosolic redox status in crypt base in the distal colon. (A) qRT-PCR for *Noxs* expression in proximal colons and distal colons. Values were normalized to *Rpl13a*. Each dot indicates one mouse. (B and C) RNAScope images (B) and quantification of relative *Nox1* mRNA intensity (C) of vertical sections of freshly frozen proximal colons and distal colons, probed for *Nox1* mRNA in red, counterstained with Hoechst in blue. Each dot indicates one ROI (3-4 ROI/mouse, n=3 mice). (D) RNAScope images of vertical sections of freshly frozen proximal colons and distal colons, probed for *Nox1* mRNA (green) and *Lgr5* mRNA (red) (n=3 mice). (E and F) Representative 405/488-nm ratio image (E) and quantification of 405/488-nm ratios (F) of vertical sections from freshly frozen distal colons from R26-RoGFP2-Orp1^Villin^ mice. Each dot indicates one crypt (n=3 mice). (G and H) Schematic of the generation of NOX1^KO^; R26-RoGFP2-Orp1^Villin^ mice (left) and Representative 405/488-nm ratio images (right) and quantification of 405/488-nm ratios (H) of distal colon explant from (G) mice. Each dot indicates one ROI (3-4 ROI/mouse, 3 mice per group). (I and J) Flow cytometry plot (I) and quantification of 405/488-nm ratios (J) from distal colon-derived colonoids of (G) mice. Each dot indicates one cell (n=3 mice). (K and L) Representative 405/488-nm ratio images (K) and quantification of 405/488-nm ratios (L) from distal colon-derived colonoids of (G) mice. Each dot indicates one colonoid (n=3 mice). Experiments were performed no less than 3 times. Scale bars, as indicated. Data in (A), (C), (F), (H), (J) and (L) are presented as mean ± SEM. Statistical significance was determined using two-tailed unpaired Student’s t tests in (C), (F), (H), (J) and (L), and Two-way ANOVA in (A); ns, not significant; ****p < 0.0001. See also Figure S1.

### NOX1-derived ROS regulates the balance of self-renewal and differentiation in the distal colon in homeostasis and during regeneration

Given the very specific expression of *Nox1* in crypt base epithelial cells along with *Lgr5*, we hypothesized that NOX1 may play a role in regulating epithelial cellular homeostasis in the distal colon. Assessment of colonic tissues in NOX1^KO^ mice indicated shorter total colon length, distal crypt depth and reduced epithelial cellularity per crypt compared to cage-matched littermate WT mice (Figures S2A and 2A-C), suggesting that loss of NOX1 is associated with altered epithelial homeostasis at baseline. Measurement of actively cycling epithelial cells via BrdU incorporation experiment showed that NOX1^KO^ mice have reduced proliferation in the distal colon (Figures 2D and 2E), but not in the proximal colon (Figures S2B and S2C), indicating a region-specific effect of NOX1. Consistent with previous reports,^22^ loss of NOX1 leads to increased proportions of MUC2^+^ goblet cells. In addition, we also found increases in other secretory lineage cells including REG4^+^ deep crypt secretory cells and CHGA^+^ enteroendocrine cells in NOX1^KO^ mice (Figures S2D-G). To assess the role of NOX1 in active regeneration of the epithelial compartment, we utilized a fasting-refeeding model (Figure S2H),^23^ to avoid from confounding tissue damage and overt inflammation seen in mucosal injury-based models.^24^ In WT mice we observed that the proportions of Ki67^+^ cycling IECs were strongly decreased upon fasting and subsequently increased above upon refeeding (Figures S2I and S2J) indicating active regeneration.^23^ Interestingly, *Nox1* expression and cellular H_2_O_2_ levels (Figures 2F, 2G and S2K) in crypt cells followed a very similar and parallel pattern of changes to Ki67 implying a role for NOX1 function in IEC regeneration upon refeeding. Further evidence for this was provided by NOX1 deficient mice which showed comparable body weight changes (Figure S2L) but impaired regeneration upon refeeding (Figures 2G and 2H).

**Figure 2.**
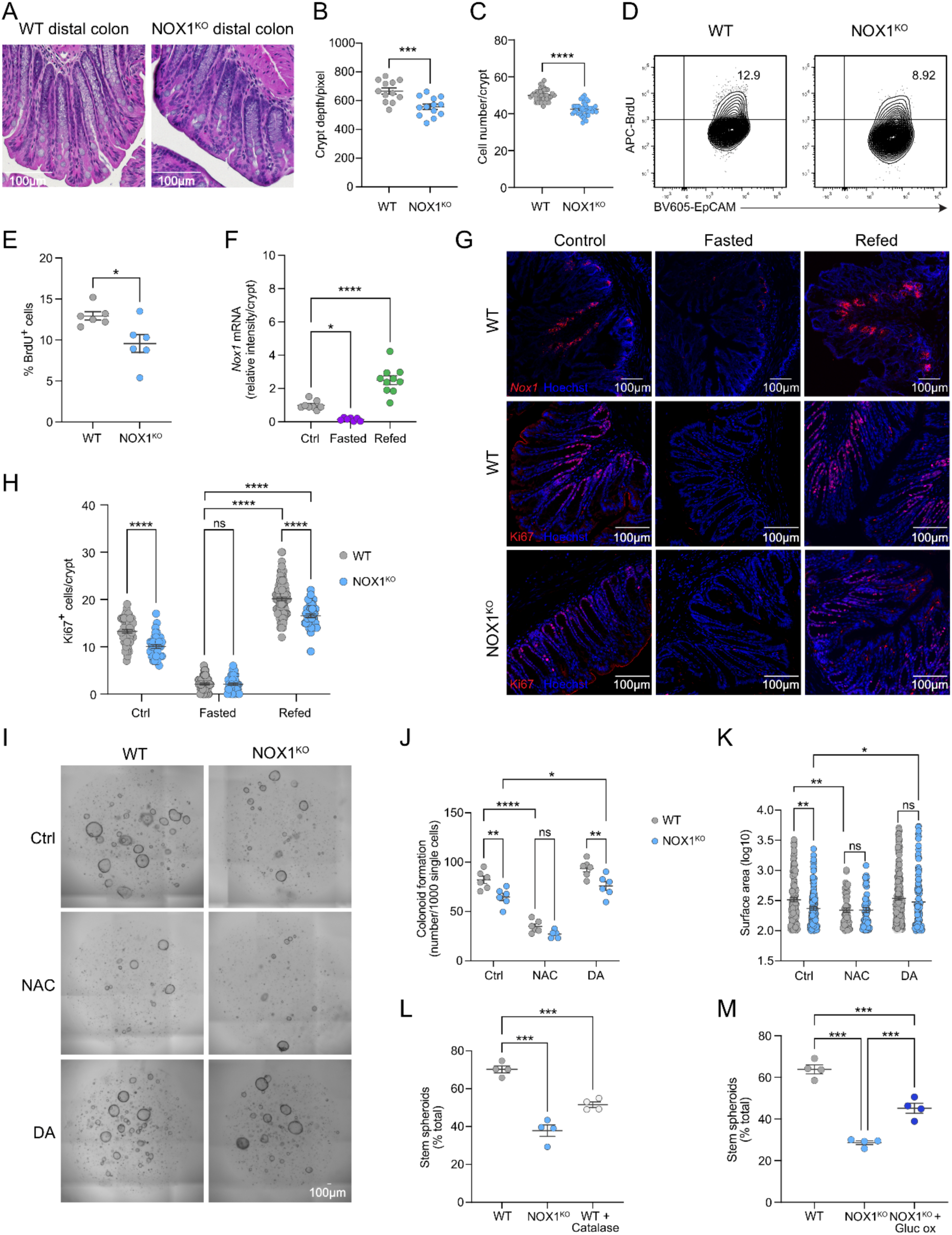
NOX1-derived ROS regulates the balance of self-renewal and differentiation in the distal colon in homeostasis and during regeneration. (A-C) H&E staining images (A), quantification of crypt depths (B) and cell number per crypt (C) of distal colons from WT and NOX1^KO^ littermates under baseline. Each dot indicates one ROI in (B) and one crypt in (C) (3-4 ROI/mouse, 3 mice per group). (D and E) Flow cytometry plot (D) and quantification (E) of BrdU incorporation in intestinal epithelial cells (Live/dead^-^CD45^-^EpCam^+^) from distal colons of WT and NOX1^KO^ littermates under baseline. Each dot indicates one mouse. (F-H) Representative fluorescence images (G) and quantifications (F) of *Nox1* mRNA intensity, quantifications (H) of Ki67^+^ cells per crypt of distal colons from WT and NOX1^KO^ littermates under fasting and refeeding. Each dot indicates one ROI (3-4 ROI/mouse, n=4-5 mice per group). (I-K) Bright field images (I), colonoids formation numbers (J) and surface area (K) of colonoids generated from isolated crypts in distal colons of WT and NOX1^KO^ littermates, treated with or without 1 mM NAC, or 10 μM diamide separately. Each dot indicates one biological replicate in (J) and one organoid in (K) (n=3 mice per group). (L) Percentage stem colonoids (spheroid) in WT and NOX1^KO^ three days after plating and in WT following incubation with 200U catalase. Each dot indicates… (M) Percentage stem colonoids (spheroid) in WT and NOX1^KO^ three days after plating and in NOX^KO^ after addition of glucose oxidase (2000U) and 20mM glucose. Experiments were performed no less than 3 times. Scale bars: 100 μm. Data (B), (C), (E), (F), (H), (J), (K), (L) and (M) presented as mean ± SEM. Statistical significance was determined using two-tailed unpaired Student’s t tests in (B), (C) and (E), One-way ANOVA in (F), (L) and (M), Two-way ANOVA in (H), (J) and (K); ns, not significant; *p < 0.05, **p < 0.01, ***p < 0.001 and ****p < 0.0001. See also Figure S2 and S3.

To understand whether NOX1 mediated changes were related to cellular redox balance we turned to stem-cell derived colonoids, which allows targeted manipulation of the cellular environment and when grown in high-Wnt medium are at baseline an *in-vitro* regeneration model.^25^ Generation of colonoids from the distal colon resulted in decreased secondary organoid formation capacity in NOX1^KO^ compared to WT, as well as reduced surface area. The NOX1 deficient cellular phenotype was recapitulated in WT colonoids by reducing cellular ROS and increasing relative cellular reductive capacity with n-acetylcysteine (NAC) treatment, and conversely increasing cellular oxidation capacity using diamide partially rescued NOX1^KO^ colonoids, suggesting that NOX1-derived ROS modulates cycling IEC self-renewal (Figures 2I-K). Morphologically, NOX1^KO^ colonoids budded earlier and more extensively than WT, which was recapitulated by treatment with the ROS scavenger catalase in WT colonoids (Figure 2L). Increased steady state environmental H_2_O_2_ generated by addition of exogenous glucose oxidase partially rescued accelerated IEC differentiation in NOX1^KO^ colonoids as measured by cyst-like stem versus budded differentiated colonoids (Figure 2M). Analysis of specific intestinal epithelial cell markers in colonoids showed higher levels of secretory cell lineage markers and absorptive lineage markers, but reduced *Ki67* expression (Figure S3A). Immunofluorescence to identify mature cell types showed higher proportions of committed IECs with reduced cycling cells in NOX1^KO^ colonoids (Figures S3B-G), consistent with the findings observed in distal colonic tissues. Taken together, these data suggest that NOX1-mediated ROS production and consequent alteration of cellular redox balance modulates the balance of epithelial self-renewal and differentiation in the distal colon.

### The intestinal stem cell-intrinsic function of NOX1 regulates self-renewal and cell fate in the distal colon

We next investigated whether NOX1-dependent regulation of the balance of IEC proliferation and differentiation were due to changes in the ISC proportions and/or ISC function. We found *Lgr5* expression levels were similar between WT and NOX1^KO^ mice (Figures S4A and 3A). To further evaluate the question of a functional or proportional role in ISCs, we generated NOX1^KO^ Lgr5 reporter mice (NOX1^KO^; Lgr5^creERT2/+^, Figure S4B) to compare with WT; Lgr5^creERT2/+^ mice. Proportions of Lgr5-EGFP^+^ ISCs were similar between WT and NOX1^KO^ mice in both the distal and proximal colons (Figures 3B and S4C). However, we found that NOX1 deficient distal colonic ISCs exhibited reduced BrdU incorporation in steady state conditions with failed active regeneration upon refeeding (Figures 3C and 3D). In contrast, and providing further evidence of a region-specific role, loss of NOX1 had no impact on the proportion of active cycling ISCs in the proximal colon in both the steady state and during regeneration (Figures S4D and S4E). To further investigate the cell-intrinsic function of NOX1, ISCs were isolated from WT; Lgr5^creERT2/+^ and NOX1^KO^; Lgr5^creERT2/+^ mice separately and plated to generate colonoids. Distal colonic NOX1 deficient ISCs showed impaired primary organoid formation capacity compared to WT ISCs (Figures 3E and 3F). Consistent with our tissue data, ISCs isolated from the proximal colon displayed similar primary organoid formation capacity in both WT and NOX1^KO^ mice (Figures S4F and S4G). ISC cell fate is thought to follow a neutral drift model, whereby ISCs undergo stochastic asymmetric division with either self-renewal to maintain stemness, or generation of committed cell progenitor.^26^ To explore whether loss of NOX1 function acts to alter this cell fate decision process, we employed *in vivo* genetic lineage tracing to track ISC cell progeny. WT;Lgr5^tdTomato^ and NOX1^KO^;Lgr5^tdTomato^ mice were generated to enable ISC cell-tracing following tamoxifen induction (Figure 3G). After induction, committed IECs from ISCs are tdtomato^+^ while daughter ISCs are both EGFP^+^ and tdtomato^+^. In the distal colon, loss of NOX1 resulted in reduced accumulation of ISC-derived tdtomato^+^ cells (Figures 3H and 3I) and a significantly smaller proportion of EGFP^+^ tdtomato^+^ daughter ISCs (Figures 3J and 3K). Loss of NOX1 however had no effect on the ISC fate decisions in the proximal colon (Figures S4H-K). These results point to a highly location specific role for NOX1 dependent ROS in regulating ISC fate decisions in the distal colon.

**Figure 3.**
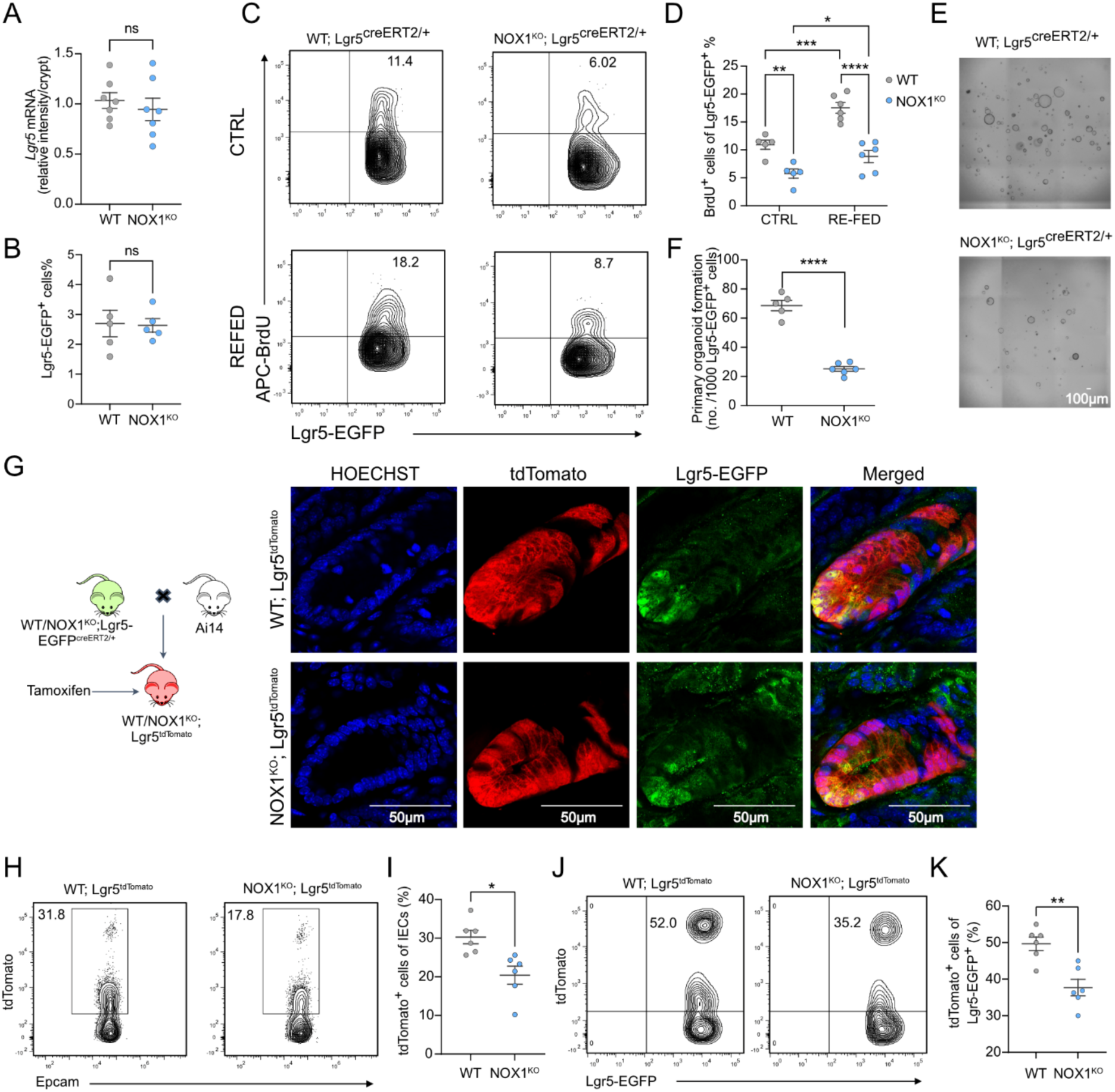
The cell-intrinsic function of NOX1 regulates ISC self-renewal and cell fate in the distal colon. (A) Quantification of *Lgr5* mRNA intensity in Figure S4A from freshly frozen distal colons. Each dot indicates one ROI (2-3 ROI/mouse, n=3 mice). (B) Proportions of Lgr5-EGFP^+^ cells (gated from Live/dead^-^CD45^-^EpCam^+^) from distal colons of WT; Lgr5^creERT2+^ and NOX1^KO^; Lgr5^creERT2+^ naïve mice. Each dot indicates one mouse. (C and D) Flow cytometry plot (C) and proportions (D) of BrdU^+^ Lgr5-EGFP^+^ cells (gated from Live/dead^-^CD45^-^EpCam^+^Lgr5-EGFP^+^) from distal colons of WT; Lgr5^creERT2+^ and NOX1^KO^; Lgr5^creERT2+^ mice under fasting and refeeding. Each dot indicates one mouse. (E and F) Bright field images (E) and quantifications (F) of primary organoid formation per 1000 sorted Lgr5-EGFP^+^ cells of distal colons from either WT; Lgr5^creERT2+^ or NOX1^KO^; Lgr5^creERT2+^ mice on day 5. Each dot indicates one biological replicate (n=2 mice). (G) Generation of lineage tracing mice WT; Lgr5^tdTomato^ and NOX1^KO^; Lgr5^tdTomato^ mice (left) and representative fluorescence images of freshly frozen distal colon sections from mice sacrificed 5 days after tamoxifen (20mg/kg) injection (right), immunolabelled for tdTomato and EGFP (n=3 mice). (H and I) Flow cytometry plot (H) and quantification (I) of total tdTomato^+^ cells (gated from Live/dead^-^CD45^-^EpCam^+^) from distal colons of (G) mice sacrificed 2 days after tamoxifen (20 mg/kg) injection. Each dot indicates one mouse. (J and K) Flow cytometry plot (J) and quantification (K) of tdTomato^+^Lgr5-EGFP^+^ cells (gated from Live/dead^-^CD45^-^EpCam^+^ Lgr5-EGFP^+^) from distal colons of (G) mice sacrificed 2 days after tamoxifen (20mg/kg) injection. Each dot indicates one mouse. Experiments were performed no less than 3 times. Scale bars, as indicated. Data (A), (B), (D), (F), (I) and (K) are presented as mean ± SEM. Statistical significance was determined using two-tailed unpaired Student’s t tests in (A), (B), (F), (I) and (K), Two-way ANOVA in (D); ns, not significant; *p < 0.05, **p < 0.01, ***p < 0.001 and ****p < 0.0001. See also Figure S4.

### Functional NOX1 regulates cell cycle entry in distal colonic ISCs and transient amplifying cells

To understand at what stage NOX1 regulates the process of ISC fate progression and capture the transcriptional signatures involved, we performed single-cell RNA sequencing (scRNA-seq) of EpCAM^+^CD45^-^ IECs from both proximal and distal colons in NOX1^KO^ and littermate WT control mice. Louvain-clustering of the resulting single-cell expression profiles revealed 17 cell-type /state clusters in distal colon (Figures 4A, S5A and S5B). This included populations of Lgr5^+^ ISCs and neighboring actively cycling transient amplifying cells (TAs). Broad clustering by genotype in distal colonic IECs showed a small shift in cluster distribution between WT and NOX1^KO^ IECs, in contrast to the proximal colon which showed overlapping distributions (Figures 4B and S5C-E). This was reflected in a shift in overall cell type/state distribution providing additional evidence for a distal colon specific role (Figures S5D, S5E and 4C). Consistent with our immunofluorescence data, NOX1^KO^ distal IECs showed increased relative abundance of secretory progenitors, secretory lineages and several sub-types of committed IECs (Figure S5E). To trace when NOX1-dependent ROS regulates the progression of ISCs we carried out trajectory analysis of our single cell data. Figure 4D shows lineage trajectories using Lgr5+ ISCs as an origin across increasing pseudotime. Focusing on the early transition from ISC to TA cells we found two distinct cell populations (Figure 4E-G) characterized by pseudotime (early and late). These populations include cells that are either Lgr5^high^Ki67^+^ (early) or Lgr5^low^Ki67^+^ (late) (Figure 4G-I) and therefore represent cells at the point of transition between ISC self-renewal and progenitor lineage commitment. A critical factor in this transition is efficient progression through cell cycle allowing stem cells to maintain the balance between self-renewal and differentiation.^27^ To identify specific transcriptional signatures during this transitional state we interrogated the relative expression of key transcription factors involved in regulating cell cycle progression (Figure S5F) and compared these between WT and NOX1^KO^ cells in both early and late TA populations. We found a distinct pattern of expression in NOX1^KO^ with reduced expression of the cell cycle transcription factor E2F1, one of several key factors for cell cycle progression from G1 phase to S phase, in conjunction with enhanced expression of E2F5, which regulates G0 phase arrest (Figures 4J and S5F).^28^ To investigate compromised cell cycle progression in NOX1^KO^ cycling cells, we carried out cell cycle analysis in colonoids. Here we found that NOX1 deficiency results in more cells arrested in G0/G1 phase and reduced S phase versus WT (Figures 4K and 4L). Taken together, this suggests that in the distal colon NOX1-derived ROS enhances progression through cell cycle to maintain the balance of self-renewal and differentiation in cycling IECs.

**Figure 4.**
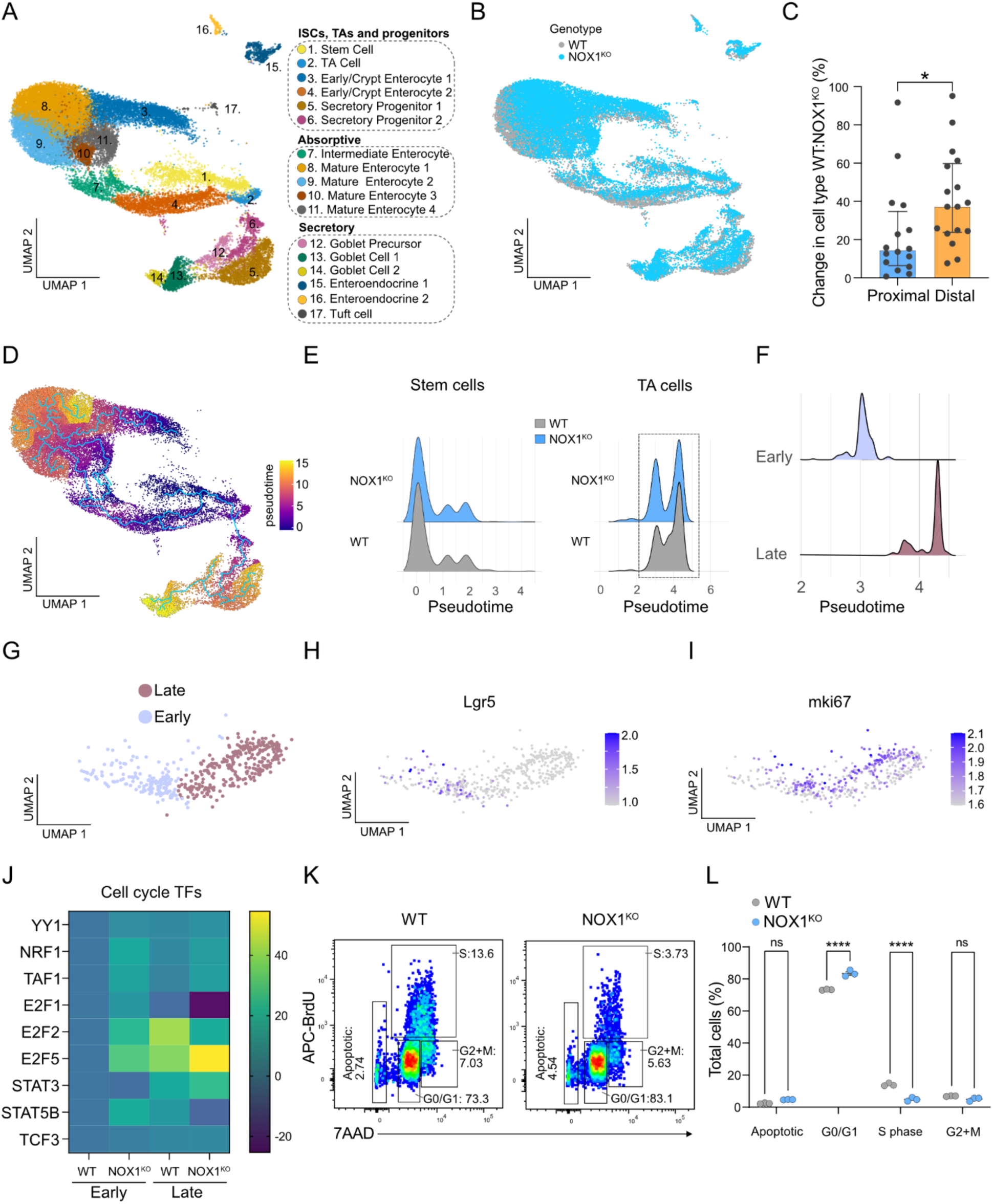
Functional NOX1 regulates cell cycle entry in distal colonic ISCs and transient amplifying cells. (A) Uniform Manifold Approximation and Projection (UMAP) of distal epithelial cell clusters. (B) Uniform Manifold Approximation and Projection (UMAP) for distal colonic epithelial cells colored by genotype-WT (grey) and NOX1^KO^ (blue). (C) Median percentage change in cell-type abundance between WT and NOX1^KO^ in distal and proximal colon (median ± SD). (D) Pseudotime trajectory analysis (Monocle3) with Uniform Manifold Approximation and Projection (UMAP) for distal colonic epithelial cells with pseudotime trajectory across epithelial clusters. (E) Cell proportions across pseudotime for stem cell and transit amplifying cell clusters in WT and NOX1^KO^. (F) Total cell proportions for TA cell cluster binned by pseudotime showing two populations (early and late). (G) Individual cells within TA cell cluster labelled by pseudotime population. (H) Relative *Lgr5* expression in TA cells. (I) Relative *mki67* expression in TA cells. (J) Heatmap of cell-cycle associated transcription factors in WT and NOX1KO TA cells in both early and late pseudotime populations. (K and L) Representative flow plots (K) and quantification (L) of cell cycle analysis in WT and NOX1^KO^ colonoids. Experiments in (K) and (L) were performed no less than 3 times. Data (C) is presented as median ± SD, and (L) is presented as mean ± SEM. Statistical significance was determined using two-tailed unpaired Student’s t tests in (C), Two-way ANOVA in (L); ns, not significant; *p < 0.05 and ****p < 0.0001. See also Figure S5.

### Loss of functional NOX1 leads to rewiring of ISC metabolism through regulation of IDH1 enzyme activity

To explore the underlying mechanisms of how NOX1-dependent ROS alters ISC function we carried out redox proteomics using the oxidative isotope-coded affinity tags (OxICAT) method^29^ to assess NOX1-dependent oxidative post-translational modifications on protein cysteine residues.^30^ Using stem-like colonoids we analyzed the relative oxidation of protein cysteines in WT versus NOX1^KO^, identifying 63 proteins with differentially oxidized cysteine residues (Figure 5A). Gene set enrichment analysis of this data using gene ontology (GO) indicated that the most significant pathway changed in NOX1^KO^ cells was inhibition of glycolysis (Figure 5B). Several studies have shown that cell metabolism and switching of energy production pathways can dictate intestinal stem cell fate decision in variety of contexts.^6,8–10,31–33^ ISC self-renewal has been linked with reliance on glycolysis while lineage commitment and differentiation with oxidative phosphorylation (OXPHOS).^34^ To initially assess metabolic alterations in the setting of NOX1 deficiency, we looked at expression of key glycolytic genes. We found significantly reduced expression of genes associated with glycolytic metabolism in NOX1 deficient cycling IECs (Figure 5C). To directly investigate cellular metabolism we carried out intracellular metabolic status analysis.^35^ We found that NOX1^KO^ IECs had lower extracellular acidification rate (ECAR) compared to WT, indicating reduced utilization of glycolytic metabolism (Figures 5D-F). Conversely, NOX1^KO^ IECs displayed enhanced oxidative phosphorylation, as measured by a significantly increased oxygen consumption rate (OCR) (Figures S6A-D), higher mitochondrial DNA copy (Figure S6E) and enhanced mitochondrial membrane potential (Figures S6F and S6G). Network analysis of our redox proteomic data revealed that altered carbon metabolism and glycolysis was one of the main features of differentially oxidized proteins (Figure 5G). One of the most differentially altered proteins and a central node in the NOX1-dependent glycolysis network was the cytoplasmic isocitrate dehydrogenase 1 (IDH1). IDH1 is a key cellular NADP^+^-dependent metabolic enzyme which reversibly catalyzes the oxidative decarboxylation of isocitrate to non-mitochondrial alpha-ketoglutarate (αKG), an important metabolite regulator of cell metabolism and proliferation.^36^ Measurement of IDH1 enzyme activity in colonoids revealed higher IDH1 enzyme activity in NOX1 deficiency (Figure 5H). Consistent with this, cellular αKG levels from either NOX1^KO^ colonoids or distal colonic tissues were higher than in WT equivalents (Figure 5I). Given previous reports that supplementation of αKG can promote differentiation and suppresses proliferation in patient-derived colon tumor organoids,^37,38^ we hypothesized that manipulation of cellular αKG levels could either rescue or phenocopy the proliferation status of NOX1^KO^ or WT cycling IECs respectively. We found cell-permeable αKG treatment could suppress cell cycling and proliferation, recapitulating the phenotype induced by NOX1 deficiency (Figures 5J and 5K). Conversely, administration of a wild type IDH1 inhibitor (compound 13)^39^ was able to restore the reduced cycling and proliferation observed in NOX1^KO^ colonoids (Figures 5J and 5K). Taken together, these results suggest that NOX1 alters the metabolic status of distal colonic cycling cells and acts via regulation of IDH1 enzyme activity to modulate ISCs function.

**Figure 5.**
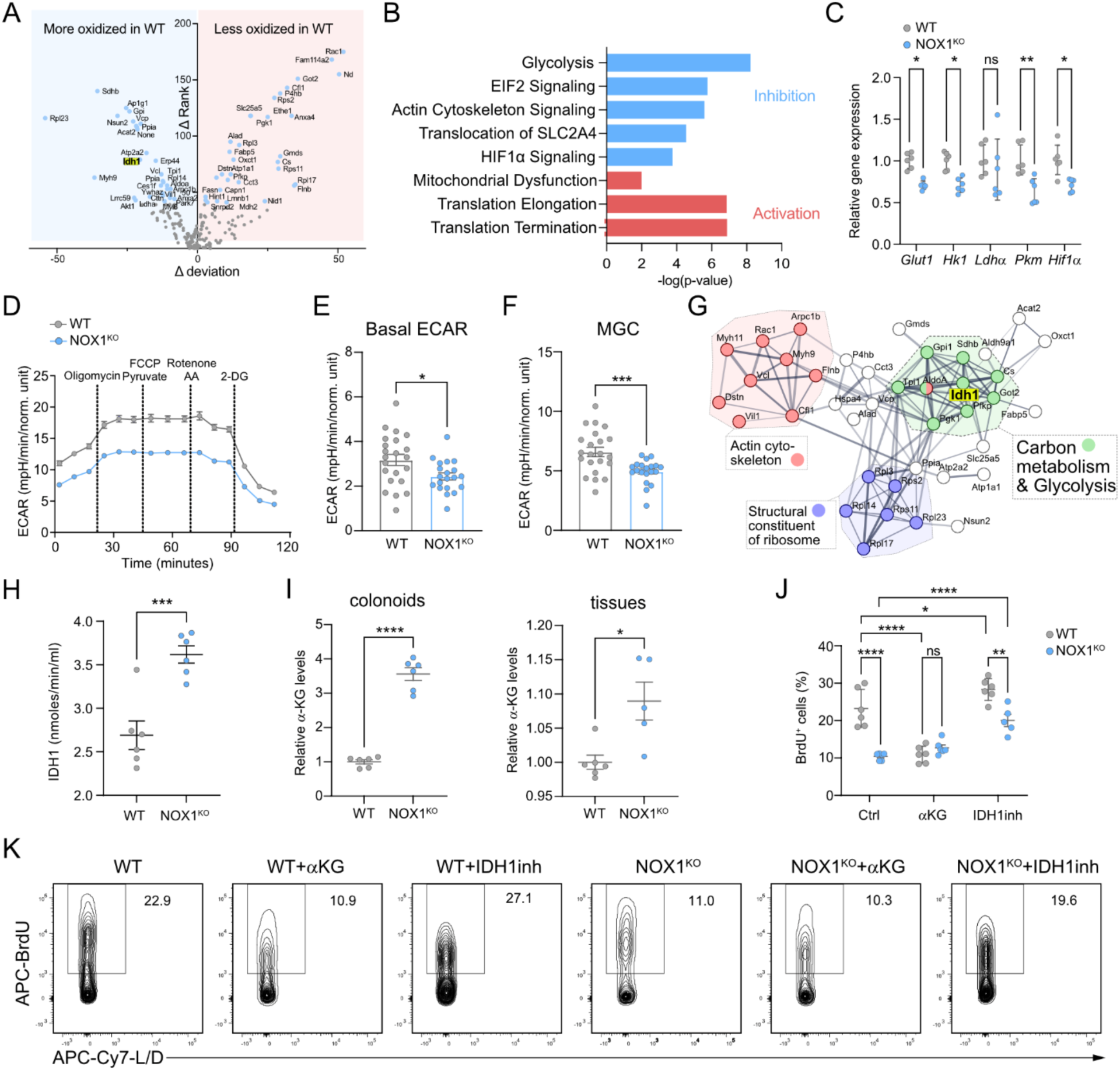
Loss of functional NOX1 leads to rewiring of ISC metabolism through regulation of IDH1 enzyme activity. (A) Volcano plot of redox proteomics highlights 63 differentially oxidized cysteine residues of proteins (change in deviation of oxidation versus change in relative rank; blue dots represent significantly altered proteins) in NOX1^KO^ colonoids versus WT control colonoids. (B) Ingenuity Pathway Analysis of significantly altered proteins from (A). (C) qRT-PCR for glycolytic gene expression in colonoids from WT and NOX1^KO^ littermates cultured in Wnt-rich WERN medium. Values were normalized to *Rpl13a*. Each dot indicates one biological replicate (n=3 mice). (D-F) Kinetic line graph (D) of extracellular acidification rate (ECAR) in 2D epithelial monolayers from WT and NOX1^KO^ littermates measured by a Seahorse Analyzer. (E) is basal ECAR and (F) is MGC bar graphs. Each dot indicates one biological replicate (n=3 mice). (G) String network analysis of (A) with enriched terms highlighted. (H) IDH1 enzyme activity assay of colonoids from distal colons of WT and NOX1^KO^ littermates (n=3 mice). (I) α-Ketoglutarate levels assay of colonoids (left) from distal colons as well as distal colon tissues (right) of WT and NOX1^KO^ littermates. Each dot indicates one biological replicate (n=3 mice) in (left) and one mouse in (right). (J and K) Flow cytometry plot (K) and quantification (J) of BrdU incorporation in colonoids from distal colons of WT and NOX1^KO^ littermates, treated with or without 3 mM α-Ketoglutarate or 2.5 mM IDH1 inhibitor (compound 13). Each dot indicates one technical replicate (n=3 mice). Experiments were performed no less than 3 times. Data (C), (D), (E), (F), (H), (I) and (J) are presented as mean ± SEM. Statistical significance was determined using two-tailed unpaired Student’s t tests in (E), (F), (H) and (I), Two-way ANOVA in (C) and (J); ns, not significant; *p < 0.05, **p < 0.01, ***p < 0.001 and ****p < 0.0001. See also Figure S6.

### The oxygen environment of the distal colonic basal crypt niche regulates NOX1 expression and function via HIF1α signaling

ISC self-renewal and differentiation is critically dependent on input by surrounding environmental niche factors.^2,40^ Our data showing that both the expression and functional effects of NOX1 are restricted to the distal colon led us to explore regionally-specific cues in the crypt base niche environment. The colon harbors the greatest microbial load within the intestine,^41^ and therefore as with other ROS-generating proteins such as Duox2,^19^ we speculated whether microbial cues may regulate the highly specific localization of *Nox1* expression. We therefore investigated *Nox1* expression in the colon of GF mice. Here we found the expression pattern of *Nox1*was identical to that seen in SPF mice, with *Nox1* highly expressed in the distal colonic crypt base, compared to proximal colon (Figures S7A-C) and suggesting against a major role for intrinsic regulation by the microbial environment.

A major niche factor that also varies along the longitudinal axis of the intestine is tissue oxygen tension, with studies suggesting that the distal colonic mucosa is a relatively more hypoxic environment relative to more proximal intestinal regions.^42,43^ Cellular responses to oxygen tension are primarily mediated by hypoxia-inducible factors (HIFs) such as HIF1α.^42,44^ To investigate this we analyzed the spatial distribution of HIF1α protein in the proximal and distal colon. In addition to increased expression within surface epithelia, stabilized HIF1α was also detected in crypt bases (Figures 6A and 6C).^45^ We found that *Hif1α* mRNA expression levels were higher (Figure S7D), and that there was significantly more stabilized HIF1α protein in crypt base cells in the distal versus proximal colon (Figures 6A and 6B). In keeping with the data on *Nox1* expression, GF mice also showed identical HIF1α expression pattern along the colon as SPF mice (Figures S7E-H), providing further evidence for an intrinsic role for hypoxia/HIF1α but not microbes. Analysis of NOX1^KO^ versus WT distal colon, showed that HIF1α was more destabilized in NOX1^KO^ tissue by both IHC and immunoblot analysis (Figures 6D-F) suggesting NOX1 function is intrinsically linked to the hypoxic niche environment.

**Figure 6.**
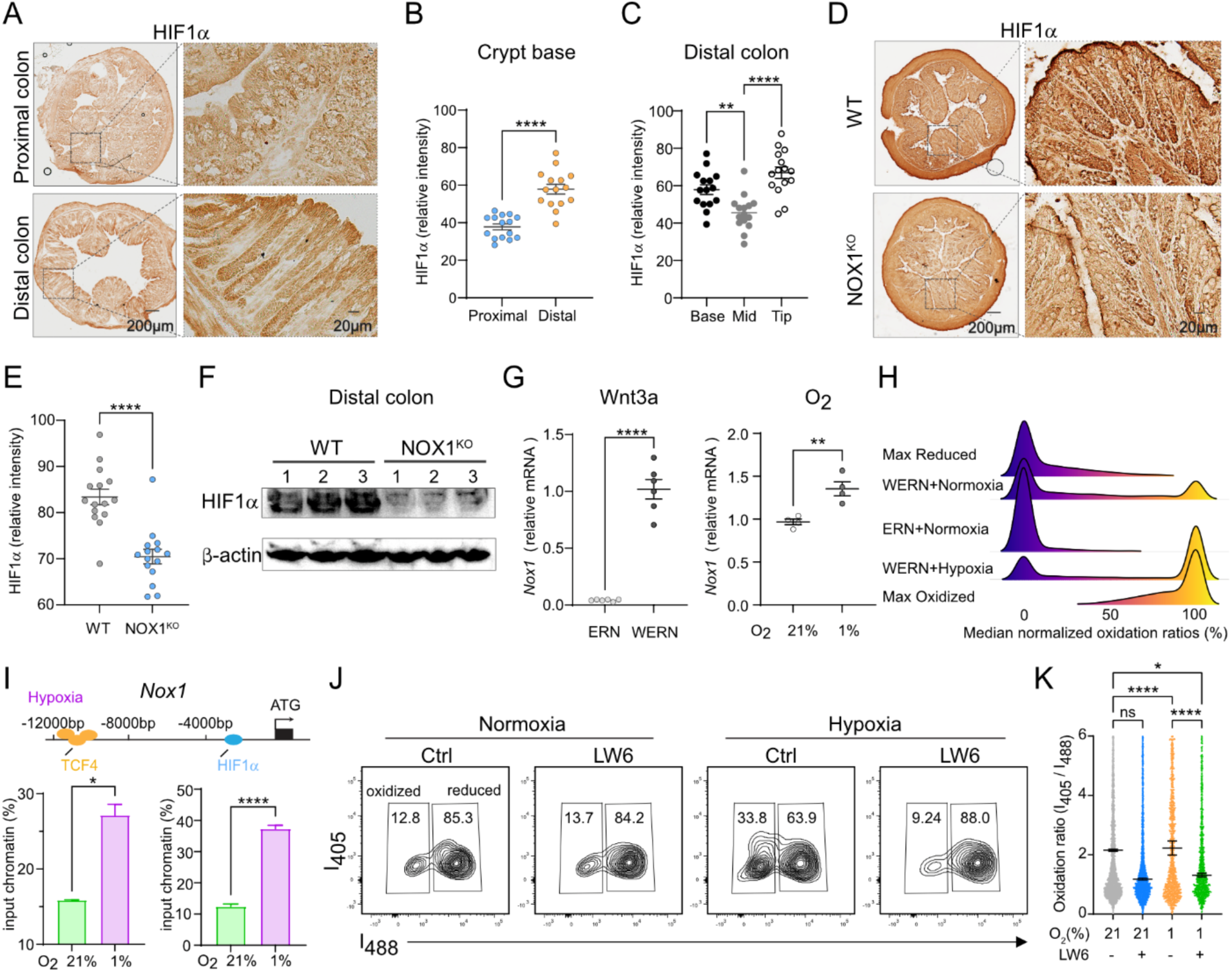
The oxygen environment of the distal colonic basal crypt niche regulates NOX1 expression and function via HIF1α signaling. (A-C) DAB analysis (A) and quantifications (B and C) of HIF1α accumulation in proximal and distal colon FFPE samples. Each spot indicates one ROI (5 ROI/mouse, n=3 mice). (D and E) DAB analysis (D) and quantification (E) of HIF1α accumulation in distal colon FFPE samples from WT and NOX1^KO^ littermates. Each dot indicates one ROI (5 ROI/mouse, n=3 mice). (E) Western blot of HIF1α protein in distal colon tissues from WT and NOX1^KO^ littermates. β-actin as the loading control (n=3 mice). (F) qRT-PCR for *Nox1* expression in colonoids from distal colons of R26-RoGFP2-Orp1^Villin^ mice cultured either in Wnt-rich WERN medium or non-Wnt ERN medium under normoxia (21% O_2_), or in Wnt-rich WERN medium under hypoxia (1% O_2_). Values were normalized to *Rpl13a*. Each dot indicates one biological replicate (n=3 mice). (G) Oxidation of roGFP in response to treatment indicated in (G). Maximum reduction (purple) and oxidation (yellow) of roGFP following additions of 2 mM Dithiothreitol (DTT) or 10 μM diamide respectively are shown as a reference. (H) Schematic (upper) of transcription factors HIF1a and TCF4 binding sites of *Nox1* gene. Primers for DNA fragments near HRE element (−2316 bp) and enhancer (−10800 bp) are used for following ChIP-qPCR. Human colonoids H514 were cultured under normoxia (21% O_2_), or hypoxia (1% O_2_) for 4 h, ChIP-qPCR was carried out with DNA fragments immunoprecipitated with anti-HIF1α (left under) or anti-TCF4 (right under) antibody separately. Values are relative to input signals. (J and K) Flow cytometry fluorescence measurements (J) and quantification (K) of 405/488-nm ratios in distal colon-derived colonoids of R26-RoGFP2-Orp1^Villin^ mice under normoxia (21% O_2_), or hypoxia (1% O_2_), treated with or without HIF1α inhibitor 10 μM LW6. Each dot indicates one cell (n=3 mice). Experiments were performed no less than 3 times. Scale bars, as indicated. Data (B), (C), (E), (G), (I) and (K) are presented as mean ± SEM. Statistical significance was determined using two-tailed unpaired Student’s t tests in (B), (E), (G) and (I), One-way ANOVA in (C), Two-way ANOVA in (K); ns, not significant; *p < 0.05, **p < 0.01 and ****p < 0.0001. See also Figure S7.

We next employed colonoids to screen the impact of a variety of niche factors on regulation of NOX1 expression and function. In addition to hypoxia, the crypt base also exhibits high concentrations of the signaling factor Wnt3a, which supports the self-renewal of ISCs.^46,47^ We exposed colonoids to niche factors present at the crypt base including Wnt3a and reduced oxygen tension versus microbial cues such as LPS and butyrate which are normally restricted to the crypt tip and surface epithelium in health.^48–50^ We found high Wnt3a and lower oxygen efficiently induced *Nox1* mRNA expression (Figures 6G). Consistent with other reports,^20,49^ butyrate reduced while LPS increased *Nox1* expression (Figure S7I). To assess function, we conducted experiments using colonoids generated from our roGFP2-Orp1 redox reporter mice. Here we found that the presence of hypoxia in the presence of high Wnt3a resulted in increased H_2_O_2_ concentrations producing a relatively oxidizing cellular environment versus either normoxia alone or normoxia along with high Wnt3a (Figure 6H), suggesting hypoxia and Wnt3a synergistically regulate NOX1 function. Consistent with the effects on expression, butyrate lowered the relative cellular oxidation level, and LPS stimulated NOX1-dependent ROS (Figure S7J).

Given the synergistic activation of NOX1 by hypoxia and Wnt3a, we investigated whether *Nox1* gene expression is directly regulated by these factors. HIF1α protein is stabilized by hypoxia and binds to hypoxia responsive elements (HREs) to drive a set of responsive genes to help cells adapt to the hypoxic environment.^44^ We therefore interrogated the promotor region of *Nox1* and located a putative HRE.^51^ Based on a database of human transcription factor targets, we also found that TCF4, a Wnt3a responsive transcription factor, has also been reported to bind around the TSS region of *Nox1*.^52^ To directly test whether *Nox1* is a hypoxia responsive gene we then performed ChIP-qPCR and found that both HIF1α as well as TCF4 bind to the regulatory regions of *Nox1* in a hypoxia dependent manner (Figure 6I). To further validate a key role of HIF1α in regulating NOX1 function we treated our hydrogen peroxide reporter colonoids with a HIF1α specific inhibitor (LW6) and assessed cellular redox balance. We found that loss of HIF1α activity under hypoxic conditions led to a relatively reduced cytosolic cellular environment confirming that HIF1α stability is required for enhanced NOX1 function under hypoxia (Figures 6J and 6K). Taken together, these data suggest that the unique hypoxic niche of the distal colonic crypt base, through the synergistic action of HIF1α and Wnt3a, allows ISCs to maintain a relatively oxidized cytosolic environment.

### Cellular redox balance orchestrates metabolic status and niche factors to control ISC cell fate

Our data showing that niche factors within the crypt base cooperate to maintain NOX1 function in the distal colon pointed towards a key role for cellular redox in regulating ISC cell fate. To explore this further we carried out bulk RNA-seq analysis in colonoids from the distal colon of NOX1^KO^ mice compared to WT littermates identifying >600 differentially expressed genes (Figure 7A). Gene set enrichment and network analysis (Figures 7B and S8A) pointed to a central role for glutathione metabolism in the transcriptional signature following loss of NOX1. Redox balance in the cell is in part set by the amount of cytosolic reducing equivalents, with glutathione (GSH) levels, quantitatively, the primary determinant. Assessment of the GSH/GSSG ratio in colonoids revealed significantly higher ratios in NOX1^KO^ colonoids (Figure 7C) both in normoxic and hypoxic conditions. Given our data showing the importance of IDH1 activity, αKG levels and HIF1α in mediating effects on metabolism and subsequently ISC proliferative capacity we investigated the effect of shifting redox balance. Manipulation of cellular redox balance towards more reduced with NAC, or more oxidized with diamide treatment, induced alterations in IDH1 enzyme activity status and αKG levels that mirrored the changes observed between WT and NOX^KO^ cells (Figures 7D and S8B). Correspondingly, under hypoxic conditions, stability of HIF1α in WT colonoids was lower in reducing cellular conditions (NAC), and similar to the administration of additional αKG.

**Figure 7.**
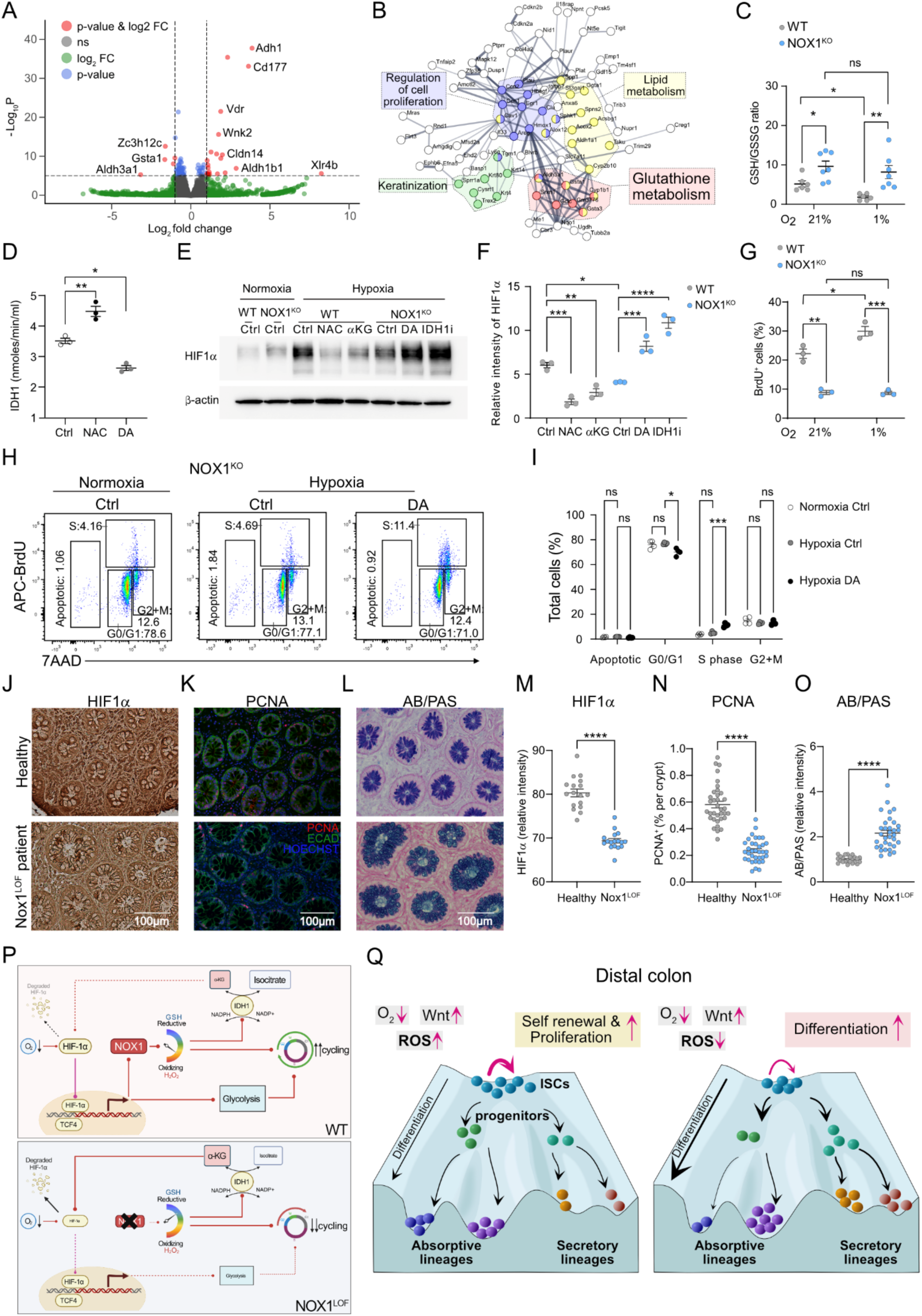
Cellular redox balance orchestrates metabolic status and niche factors to control ISC cell fate. (A) Volcano plot of bulk RNA-seq highlights 614 differentially expressed genes (DEGs, in red; log2 fold change >0.5, adjusted p < 0.05) in NOX1^KO^ colonoids versus WT control colonoids. (B) String network analysis of (A) with enriched terms highlighted. (C) GSH/GSSG ratio detection assay of colonoids from distal colons of WT and NOX1^KO^ littermates under either normoxia (21% O_2_) or hypoxia (1% O_2_) for 4 h. Each dot indicates one biological replicate (n=3 mice). (D) IDH1 enzyme activity assay of colonoids treated with or without 5 mM NAC or 10 μM diamide. Each dot indicates one biological replicate. (E and F) Representative western blot images (E) and quantification (F) of HIF1α protein in colonoids from WT and NOX1^KO^ littermates under either normoxia (21% O_2_) or hypoxia (1% O_2_) for 4 h, WT colonoids were treated with or without 5 mM NAC, 3 mM α-Ketoglutarate separately as indicated, NOX1^KO^ colonoids were treated with or without 300 μM diamide, 2.5 mM IDH1 inhibitor (compound 13). β-actin as the loading control. (G) Flow cytometry measurements of BrdU incorporation in distal colon-derived colonoids of WT and NOX1^KO^ littermates under normoxia (21% O_2_), or hypoxia (1% O_2_) for 2 h in Figure S8F. Each dot indicates one biological replicate. (H and I) Representative flow plots (H) and quantification (I) of cell cycle analysis in NOX1^KO^ colonoids under normoxia (21% O_2_), or hypoxia (1% O_2_) for 2 h, treated with or without 300 μM diamide. Each dot indicates one biological replicate. (J-O) DAB analysis (J) and quantification (M) of HIF1α intensity, fluorescence images (K) and quantifications (N) of PCNA^+^ cells, AB/PAS staining (L) and quantification of its relative intensity (O) in healthy patient and Nox1 loss-of-function patient sigmoid FFPE samples. (P) Model illustrating that rheostat-like control of cellular redox regulates IDH1 enzyme activity, αKG production to enhance hypoxia/HIF1α signaling, which in turn transcriptionally reinforces NOX1-dependent ROS and glycolytic metabolism to promote efficient cell cycle progression. (Q) Waddington model depicting the epithelial cell-state landscape in distal colon as controlled by cellular redox. Under hypoxia, high-Wnt and a relatively oxidizing redox state, the trajectory of ISCs favors self-renewal versus lineage commitment. In the absence of NOX1-dependent ROS, the relatively reductive cellular state decreases the probability of ISC self-renewal and skews the lineage hierarchy, altering proportions of absorptive and secretory IECs. Experiments were performed no less than 3 times. Scale bars: 100 μm. Data (C), (D), (F), (G), (I), (M), (N) and (O) are presented as mean ± SEM, Statistical significance was determined using two-tailed unpaired Student’s t tests in (M), (N) and (O), One-way ANOVA in (D), Two-way ANOVA in (C), (F), (G) and (I); ns, not significant; *p < 0.05, **p < 0.01, ***p < 0.001 and ****p < 0.0001. See also Figure S8.

αKG functions as the substrate of prolyl hydroxylase domain (PHD) proteins and leads to the protein hydroxylation of HIF1α.^53^ We found αKG treatment recapitulated the destabilized HIF1α seen in NOX1^KO^ colonoids (Figures 7E and 7F) and lowered NOX1-dependent ROS generation under hypoxia (Figures S7C and S7D). Conversely, in NOX1^KO^ colonoids shifting to more oxidizing conditions or inhibiting IDH1 activity increased HIF1α stability (Figures 7E and 7F). Importantly, we found that hypoxia alone, and therefore increased HIF1α, in a reducing environment (Figures 7C, 7G and S8E) was insufficient to rescue the impaired proliferation observed following loss of NOX1. This suggests that redox balance, and maintenance of sufficient oxidative equivalents through the activity of NOX1, is central to ISC function in the distal colon. To provide further evidence for this we assessed cell cycle progression following cellular redox manipulation. Shifting the cellular environment towards reduction in WT colonoids (Figures S7F and S7G) resulted in impaired transition to S phase, phenocopying loss of NOX1. Correspondingly, a shift to increased oxidation under hypoxia in NOX1^KO^ was able to rescue cell cycle progression (Figures 7H and 7I).

In the human distal colon, loss of function variants in NOX1 have been shown to be clinically important as risk allele in very early-onset inflammatory bowel disease (VEOIBD). We had previously characterized a *Nox1* missense mutation (p.N122H) that abrogated cellular ROS production, and resulted in ulcerative colitis-like VEOIBD in a patient.^54^ Patient biopsy sections showed reductions in crypt basal HIF1α stability (Figures 7J and 7M), reduced PCNA^+^ cycling cells (Figures 7K and 7N) as well as increased goblet cell proportions (Figures 7L and 7O) mirroring the changes in ISC function observed in mouse NOX1^KO^ tissue and colonoids.

## DISCUSSION

The regulation and fate of multipotent stem cells relies on a complex interplay between environment, metabolic status and cellular regulatory factors. Our studies here show that the distal colon crypt base represents a significantly different stem cell environment from more proximal intestinal regions. In the distal colon we show that the niche environment maintains NOX1-derived ROS and synergizes to alter the balance of self-renewal and differentiation in cycling stem cells in both homeostasis and active regeneration states (Figures 7P and 7Q). We propose that relative redox balance acts as a cellular rheostat whereby a shift to either a more oxidative or reductive state is central to metabolic control of the ISC cell-cycle. The hypoxic and high-Wnt niche environment at the distal colonic crypt base transcriptionally determines the unique spatial restriction of *Nox1* expression in cycling cells, contributing to relatively more oxidized cell state. As a result, lower IDH1-dependent decarboxylation of isocitrate to αKG leads to increased HIF1α stability, enhancing both NOX1 expression and glycolysis in ISCs to metabolically maintain self-renewal.^13^

Loss of NOX1 function shifts cellular redox balance towards relative reduction or ‘reductive stress’ that lowers efficiency of cell cycle entry, stem and TA cell population progression, and ultimately predisposing towards differentiation, including both secretory lineages and absorptive lineages (Figure 7Q). Our single-cell transcriptomic data uncovered alterations in the E2 transcription factor family in NOX1^KO^ cycling transit amplifying cells with lower expression of the ‘activating’ E2F1, important in promoting S-phase initiation, and elevated expression of the ‘repressor’ E2F5.^28,55^ These findings, along with data showing that manipulation of cellular ROS and IDH1 activity can mirror the changes in cell cycle seen between WT and NOX1^KO^, links together cellular redox balance and metabolism with cell-cycle efficiency in colonic ISCs. Despite increased cellular glutathione in NOX1^KO^ cells, our data does also point to the presence compensatory mechanisms with concomitant inhibition of NRF2-regulated pathways, potentially to limit reductive stress and allow some ISC proliferation.

ROS is thought to play an important role in various colonic diseases including Crohn’s disease, ulcerative colitis, pouchitis and irritable bowel syndrome.^17,18^ Several studies have established NOX1 loss of function variants in the pathogenesis of VEOIBD.^56–58^ Examination of the role of NOX1 in colitis showed that TNF-α stimulates NOX1-dependent ROS and leads to enhanced basal lymphoplasmacytosis and alteration in M-cell signatures.^59^ A few previous studies have explored the role of NOX1 in intestinal epithelium homeostasis. Denis et al showed that NOX1 deficiency decreased proliferating cell populations while increased goblet cell numbers by attenuating Wnt as well as Notch signaling pathway in colon, in keeping with our observations of a relative increase in secretory cells.^22^ In another study, NOX1-derived ROS was found to be important for IEC proliferation in distal colon through EGFR activity as suggested by *in vitro* chemical inhibition studies.^60^ In colonic cancer stem cells, NOX1-dependent mTORC1 activation via S100A9 oxidation is important for proliferation and colon cancer progression.^61^ In many of these studies direct function of NOX1 on cellular ROS were either not assessed or limited by non-quantitative redox measurements. Here, we used a genetically encoded, quantitative ratiometric cell-specific reporter *in vivo* to precisely delineate the cellular and spatial NOX1-dependent regulation of ROS and redox balance. In conjunction with genetic lineage tracing and scRNAseq analysis, this allowed careful dissection of the complex self-reinforcing mechanism that connects HIF1α-dependent hypoxic signaling to NOX1-dependent cellular ROS regulation of isocitrate dehydrogenase 1 (IDH1) in the cytosol, and maintenance of ISC function.

IDH1 is a key cytosolic metabolic enzyme that links cellular metabolism to epigenetic regulation and redox states via control of αKG levels.^62^ αKG supplementation in *Apc*^Min/+^ organoids leads to DNA hypomethylation of genes related to differentiation, attenuating Wnt signaling and driving differentiation in colorectal cancer.^37^ Studies have revealed that wild-type IDH1 is overexpressed in several types of cancers,^62^ although we did not observe any changes in *Idh1* mRNA levels in NOX1^KO^ cycling stem cells. αKG supplementation can mimic the phenotype of NOX1-loss-of-function, decreasing redox status under hypoxia and restricting cell cycle progression. IDH1 is one of the most mutated metabolic enzymes across human cancers, with mutant IDH1 enzyme exhibiting neomorphic enzymatic activity.^53^ Mutant IDH1 generates the oncometabolite 2-hydroxyglutarate (2-HG), which inhibits αKG-dependent enzymes, including histone, DNA, and RNA demethylases, metabolic enzymes, and DNA repair factors. Our data connecting NOX1-dependent ROS to IDH1 activity may provide a link between redox state and epigenetic modification in cancer progression. In addition to links to colon cancer, the role of redox balance in setting epigenetic regulation in homeostasis or regeneration after injury in colonic ISCs and the effects on the regulatory pathways outlined by our studies need to be explored further.

An interesting finding in our study was that NOX1 and HIF1α mediated effects were intrinsic to the niche environment and independent of luminal microbial input. Previous studies have outlined the importance of HIF1α in epithelial cells during inflammation and for maintaining colonic regeneration during injury.^63–65^ Our studies show significantly increased HIF1α protein in crypt base ISCs in the distal versus proximal colon and suggestive of a different oxygen tension within the distal niche environment. In this setting, HIF1α and NOX1 form a self-reinforcing circuit to maintain cellular oxidative balance and hypoxia-induced transcriptional responses including promotion of glycolysis. Several studies have shown that changes in redox-altering proteins including NOX1, DUOX2 or the combined loss of NOX isoforms, induce microbial changes that affect epithelial function and in the case of *Duox2* are directly regulated by commensal microbes.^19,66–68^ *Nox1* is localized at in the distal colonic crypt base which is a site largely shielded from microbes in homeostasis. However, during inflammation or barrier breakdown, as shown for the microbial metabolite butyrate and for other microbial products such pathogen associated molecular patterns (PAMPs), microbial or cytokine-related signaling responses are likely modulated by NOX1-dependent ROS in crypt base stem cells.

Redox balance and signaling are thought to play a role in stem cell function in a variety of tissue compartments, but most studies have focused on the deleterious effects of excessive cellular ROS or ‘oxidative stress’. In embryonic pluripotent stem cells (PSCs) and hematopoietic stem cells (HSCs), stem cells maintain very low cytosolic ROS levels in quiescence and elevated ROS levels are associated with activation and progression through differentiation, but also decreased self-renewal and increased stem cell exhaustion.^69,70^ In contrast, in neural stem cells (NSCs), increasing ROS correlates with stem cell proliferation *in vivo,* and boosts the capacity of NSCs for neurosphere formation.^71^ These and other studies suggest that the role of redox balance in stem cells depends on context, and is likely driven by the metabolic and regenerative requirements of the specific tissue environment. Adult stem cells in mucosal epithelial tissues such as the distal intestine are required to maintain constant self-renewal and proliferation to maintain the rapidly renewing epithelial barrier and are therefore likely to require a balanced redox state to enable efficient metabolism for biomass production. Our data would suggest that in the setting of physiological hypoxia, as occurs in the distal colon, the set-point for this redox balance favors a more oxidative cellular state to enable efficient cell-cycle progression.

In summary, our data suggests that, in the distal colon, a self-reinforcing metabolic-transcriptional circuit centered around maintenance of NOX1-dependent cellular redox balance, regulates ISC function, self-renewal and subsequent cell fate decisions. Our studies show how redox-dependent stem cell metabolic state switching occurs in hypoxic tissue environments, and provide a basis for understanding regeneration dynamics, barrier function and disease propensity in the distal large intestine. Further studies are needed to understand how the dynamic control of stem cell function by hypoxia and redox influence disease-driven processes that disproportionately affect the distal colon such as ulcer formation, inflammation and cancer.

### Limitations of study

Limitations of our studies that should be addressed by future work include elucidating the precise sub-cellular localization of NOX1 and the contribution of cell-cell communication by neighboring crypt base epithelial and stromal cells. A major by-product of aerobic glycolysis is lactate^72,73^, and how NOX1-dependent ROS impacts lactate metabolism and scavenging is a key question in understanding colonic ISC function. Metabolic pathways have complex interactions with compensatory changes and exactly how electron transfer occurs between the cytosol and mitochondria to maintain oxidative equivalents in the form of NAD+ in colonic epithelia also remains poorly understood.

---

**Figure S1.**
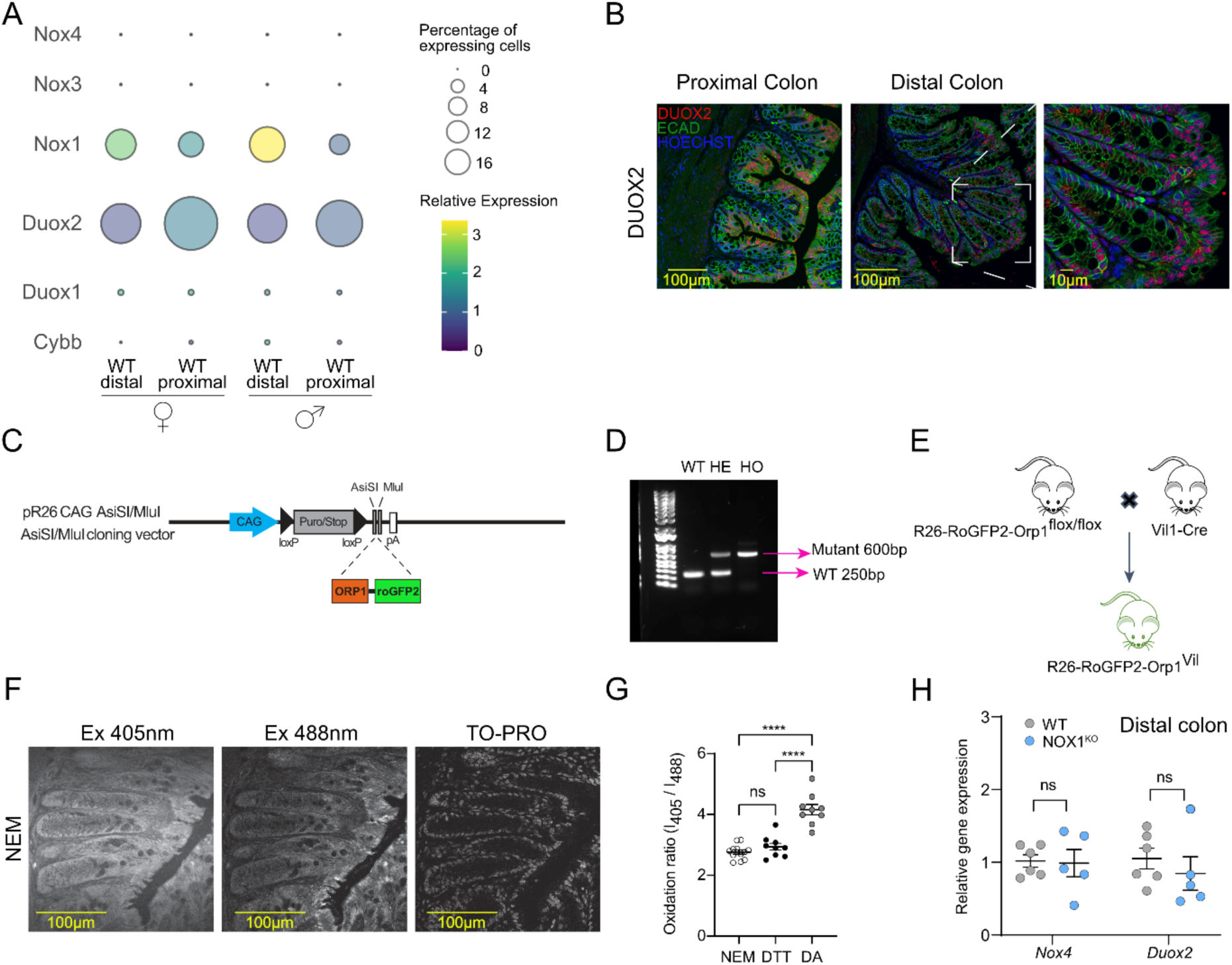
*Nox1* is highly expressed in the distal colon and regulates redox status in crypt base. (A) NADPH oxidases family mRNA expression levels from bulk RNA-seq (GSE214866) performed on purified intestinal epithelial cells from different colonic regions. (B) Fluorescence microscopy of proximal and distal colon sections from WT naïve mice immunolabelled for DUOX2 (in red) and E-cadherin (in green) (n=3 mice per group). (C) Schematic of R26-RoGFP2-Orp1^flox/flox^ mouse strain generated by cloning-free CRISPR/Cas system. pR26roGFP2Orp1 targeting vector was generated by assembling Orp1-roGFP2 to the AsiSI/MluI restriction sites in Rosa26 targeting vectors pR26 CAG AsiSI/MluI.^77^ (D) Genotype PCR of R26-RoGFP2-Orp1^flox/flox^ mouse strain. Wilde type (1^st^ lane) only has 250bp band, heterozygous (2^nd^ lane) has both 250 bp and 600 bp bands, homozygous (3^rd^ lane) has 600bp band. (E-G) Representative fluorescence images of excitation at 405 nm (left panel), 488 nm (middle panel) and far-red TO-PRO (left panel) (F) from (E) R26-RoGFP2-Orp1^Villin^ mice. Ratios of 405/488-nm from distal colon sections subjected to only 50 mM NEM or pre-treated with 20 mM DTT, 1 mM diamide (DA) separately (G). Each dot indicates one ROI (3-4 ROI/mouse, 3 mice per group). (H) qRT-PCR for *Noxs* expression in distal colons from WT and NOX1^KO^ littermates. Values were normalized to *Rpl13a*. Each dot indicates one mouse. Experiments were performed no less than 3 times. Scale bars, as indicated. Data (G) and (H) are presented as mean ± SEM. Statistical significance was determined using One-way ANOVA in (G) and Two-way ANOVA in (H); ns, not significant; ****p < 0.0001.

**Figure S2.**
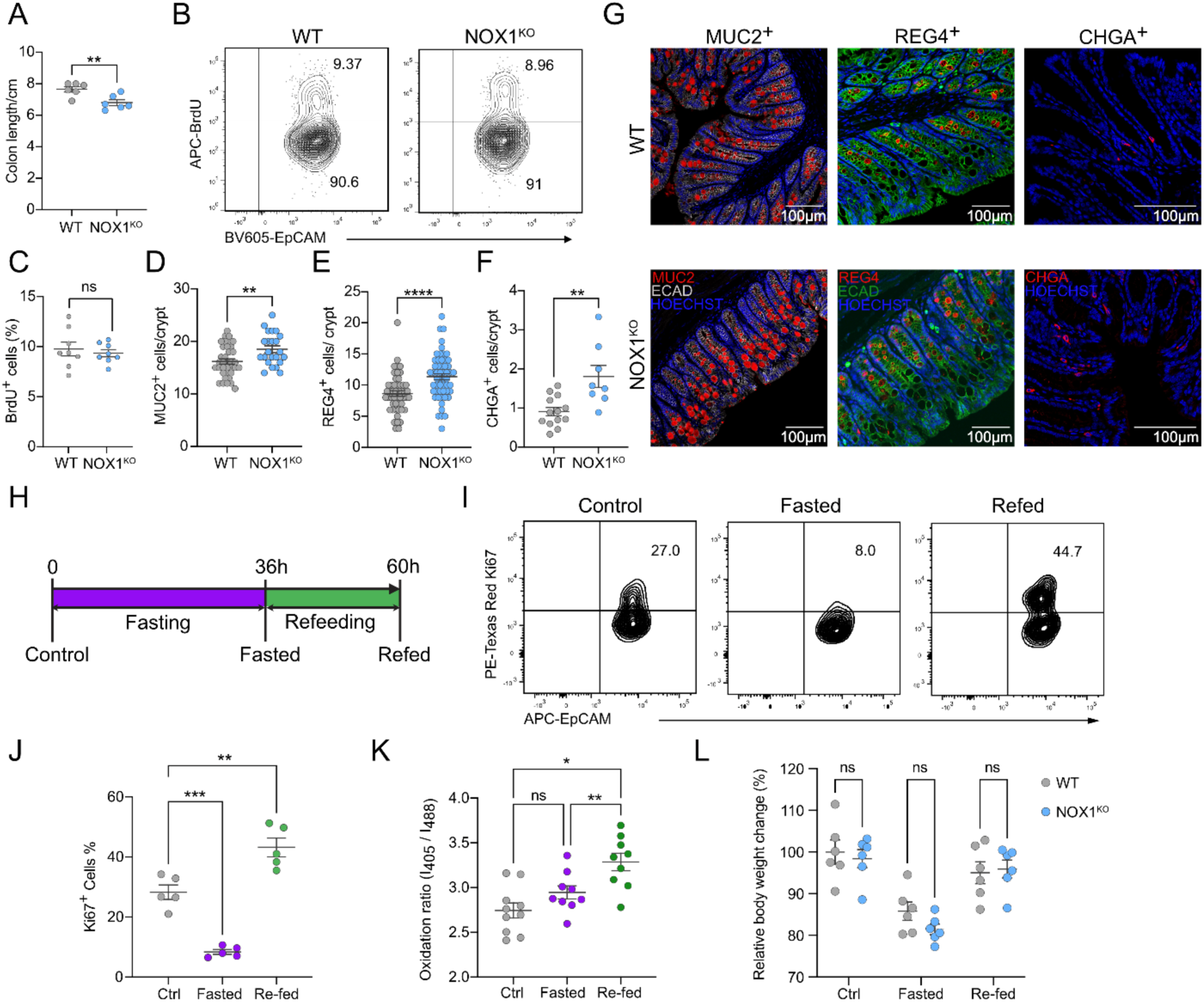
NOX1 deficiency can alter balance of proliferation and differentiation in distal colon tissues. (A) Colon lengths of WT and NOX1^KO^ littermates. Each dot indicates one mouse. (B and C) Flow cytometry plot (B) and quantification (C) of BrdU incorporation in intestinal epithelial cells (Live/dead^-^CD45^-^EpCam^+^) from proximal colons of WT and NOX1^KO^ littermates under baseline. Each dot indicates one mouse. (D-G) Representative fluorescence images (G) and quantifications (D) of MUC2^+^, (E) of REG4^+^, and (F) of CHGA^+^ cells of distal colon sections from WT and NOX1^KO^ littermates. Each dot indicates one ROI (3-4 ROI/mouse, n=3 mice per group). (H-J) Schematic of fasting-refeeding models in Figures 2G and S2I (H). Flow cytometry plot (I) and ratios (J) of Ki67^+^ intestinal epithelial cells (Live/dead^-^CD45^-^EpCam^+^) from distal colons of WT mice. Each dot indicates one mouse in (J). (K) Ratios of 405/488-nm from distal colon sections of R26-RoGFP2-Orp1^Villin^ mice in model (H). Each dot indicates one ROI (3-4 ROI/mouse, n=3 mice per group). (L) Body weight of WT and NOX1^KO^ littermates under model (H). Each dot indicates one mouse. Experiments were performed no less than 3 times. Scale bars: 100 μm. Data (A), (C), (D), (E), (F), (J), (K) and (L) presented as mean ± SEM. Statistical significance was determined using two-tailed unpaired Student’s t tests in (A), (C), (D), (E) and (F), One-way ANOVA in (J) and (K), Two-way ANOVA in (L); ns, not significant; *p < 0.05, **p < 0.01, ***p < 0.001 and ****p < 0.0001.

**Figure S3.**
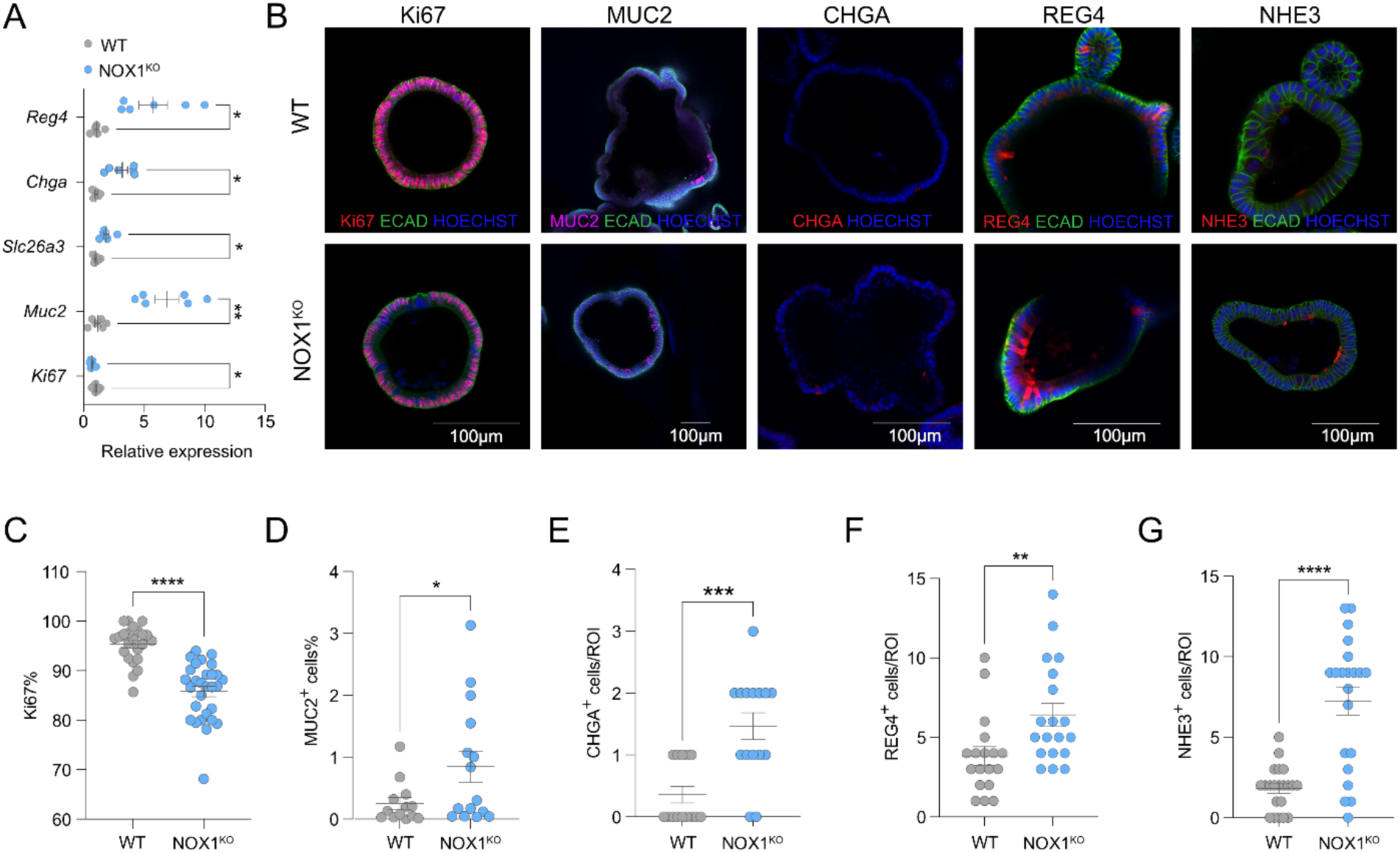
NOX1 deficiency can alter balance of proliferation and differentiation in colonoids. (A) qRT-PCR for specific intestinal epithelial cell markers expression in colonoids from WT and NOX1^KO^ littermates. Values were normalized to *Rpl13a*. Each dot indicates one biological replicate (n=3 mice). (B-G) Representative fluorescence images (B) and quantifications of Ki67^+^ (C), MUC2^+^ (D), CHGA^+^ (E), REG4^+^ (F) and NHE3^+^ (G) cells of colonoids generated from crypts of distal colons from WT and NOX1^KO^ littermates. Each dot indicates one ROI (4-7 ROI/mouse, n=3 mice per group). Experiments were performed no less than 3 times. Scale bars: 100 μm. Data (A), (C), (D), (E), (F) and (G) are presented as mean ± SEM. Statistical significance was determined using two-tailed unpaired Student’s t tests in (C), (D), (E), (F) and (G), Two-way ANOVA in (A); ns, not significant; *p < 0.05, **p < 0.01, ***p < 0.001 and ****p < 0.0001.

**Figure S4.**
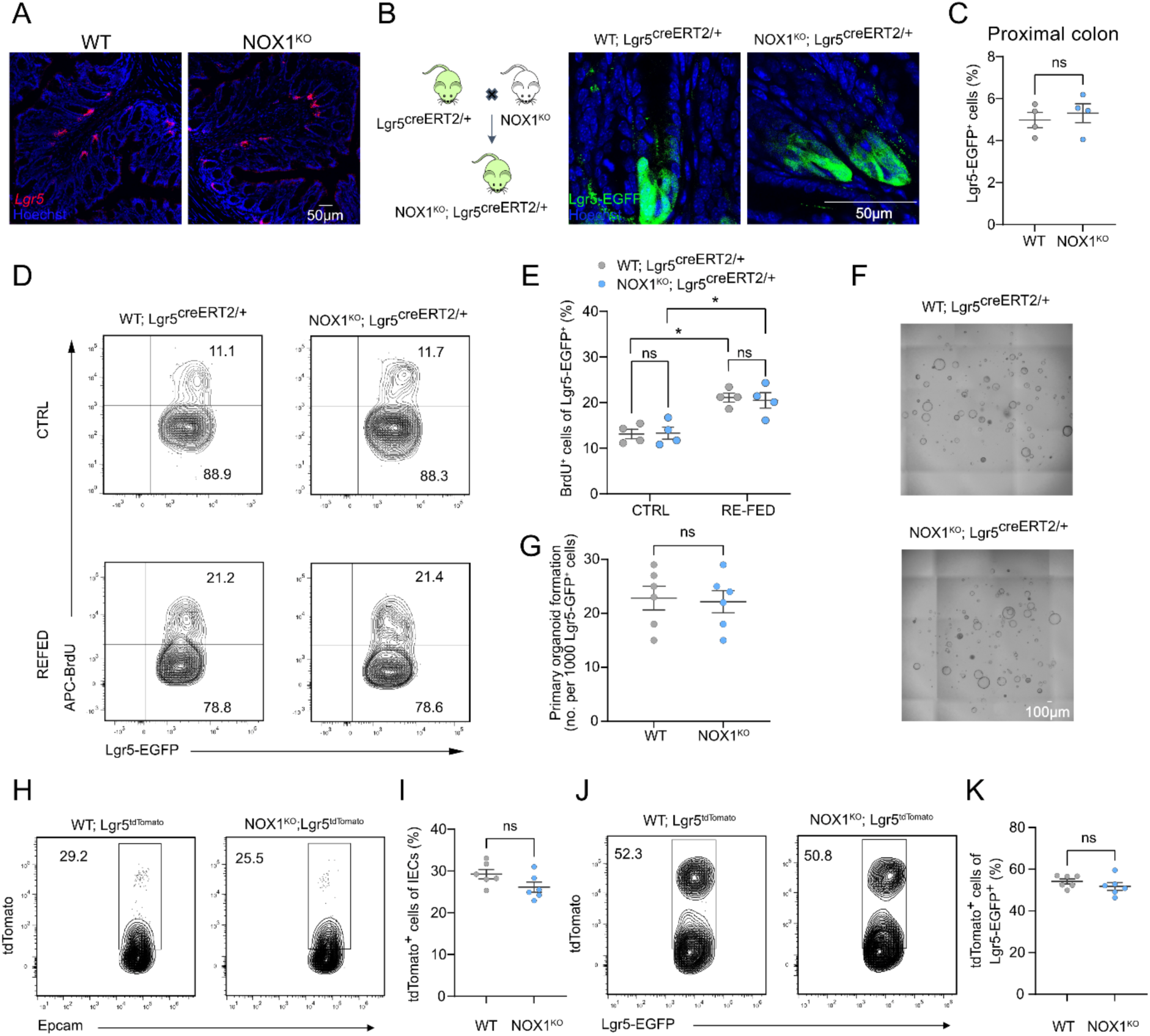
NOX1 deficiency does not alter intestinal stem cells proportions nor functions in the proximal colon. (A) Representative RNAScope images of *Lgr5* mRNA intensity from freshly frozen distal colons. See also Figure 3A. (B) Generation of NOX1^KO^; Lgr5^creERT2/+^ mice (left) and representative fluorescence images of Lgr5-EGFP^+^ cells in distal colons (right). (C) Proportions of Lgr5-EGFP^+^ cells (gated from Live/dead^-^CD45^-^EpCam^+^) from proximal colons of WT; Lgr5^creERT2+^ and NOX1^KO^; Lgr5^creERT2+^ naïve mice. Each dot indicates one mouse. (D and E) Flow cytometry plot (D) and proportions (E) of BrdU^+^ Lgr5-EGFP^+^ cells (gated from Live/dead^-^CD45^-^EpCam^+^Lgr5-EGFP^+^) from proximal colons of WT; Lgr5^creERT2+^ and NOX1^KO^; Lgr5^creERT2+^ mice under fasting and refeeding. Each dot indicates one mouse. (F and G) Bright field images (F) and quantifications (G) of primary organoid formation per 1000 sorted Lgr5-EGFP^+^ cells of proximal colons from either WT; Lgr5^creERT2+^ or NOX1^KO^; Lgr5^creERT2+^ mice on day 5. Each dot indicates one biological replicate (n=2 mice). (H and I) Flow cytometry plot (H) and quantification (I) of total tdTomato^+^ cells (gated from Live/dead^-^CD45^-^EpCam^+^) from proximal colons of (Figure 3G) mice sacrificed 2 days after tamoxifen (20 mg/kg) injection. Each dot indicates one mouse. (J and K) Flow cytometry plot (J) and quantification (K) of tdTomato^+^Lgr5-EGFP^+^ cells (gated from Live/dead^-^CD45^-^EpCam^+^ Lgr5-EGFP^+^) from proximal colons of (Figure 3G) mice sacrificed 2 days after tamoxifen (20 mg/kg) injection. Each dot indicates one mouse. Experiments were performed no less than 3 times. Scale bars, as indicated. Data (C), (E), (G), (I) and (K) are presented as mean ± SEM. Statistical significance was determined using two-tailed unpaired Student’s t tests in (C), (G), (I) and (K), Two-way ANOVA in (E); ns, not significant; *p < 0.05.

**Figure S5.**
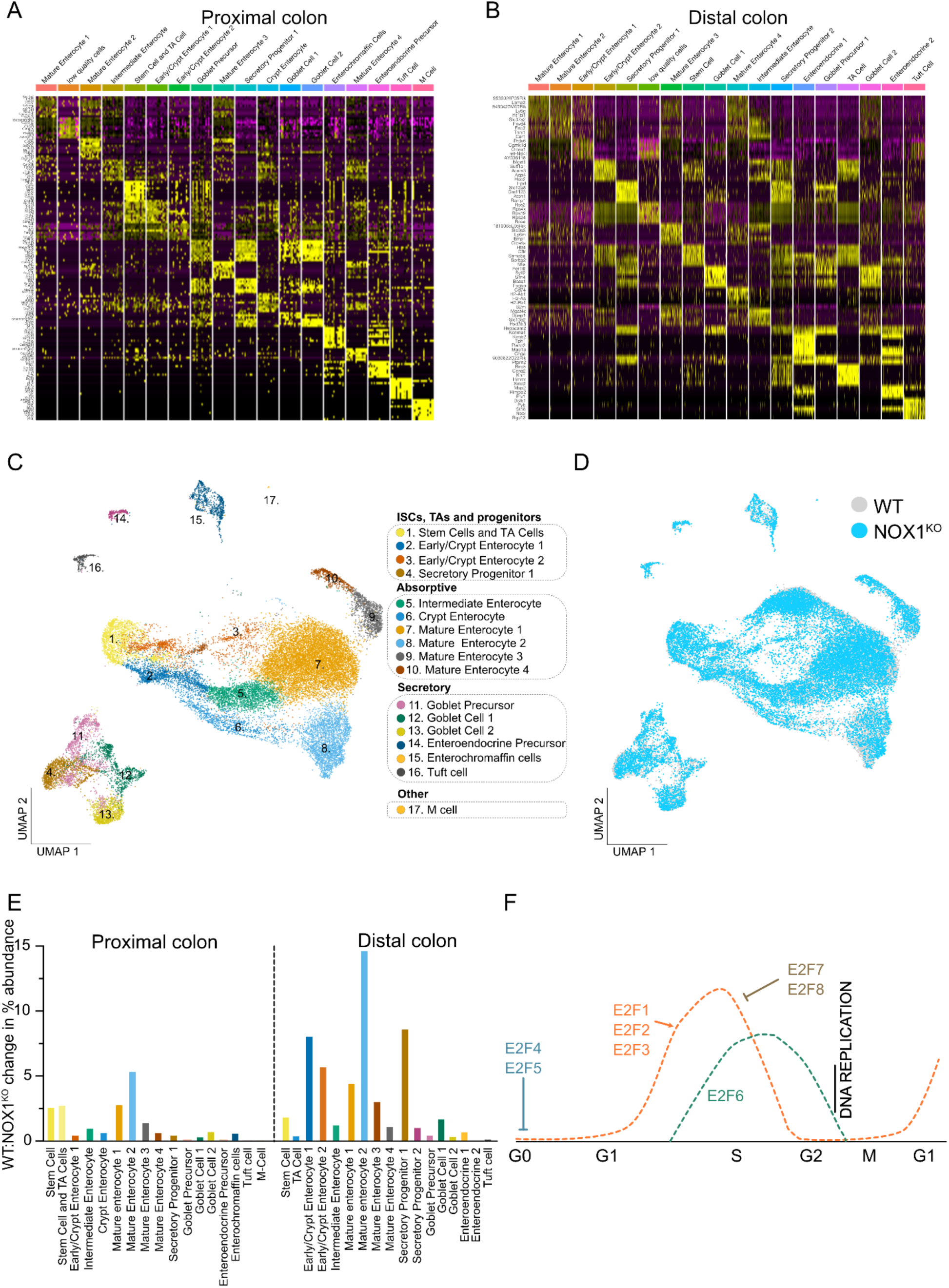
Single-cell transcriptomic analysis of proximal colon. (A) Heatmap showing marker genes defining scRNAseq cell cluster annotations in the proximal colon. (B) Heatmap showing marker genes defining scRNAseq cell cluster annotations in the distal colon. (C) Uniform Manifold Approximation and Projection (UMAP) of proximal epithelial cell clusters. (D) Uniform Manifold Approximation and Projection (UMAP) for proximal colonic epithelial cells colored by genotype -WT (grey) and NOX1^KO^ (blue). (E) Changes in cell abundance for annotated cell clusters in proximal and distal colon. (F) Schematic showing role of E2F family of transcription factors in cell cycle progression.^28^

**Figure S6.**
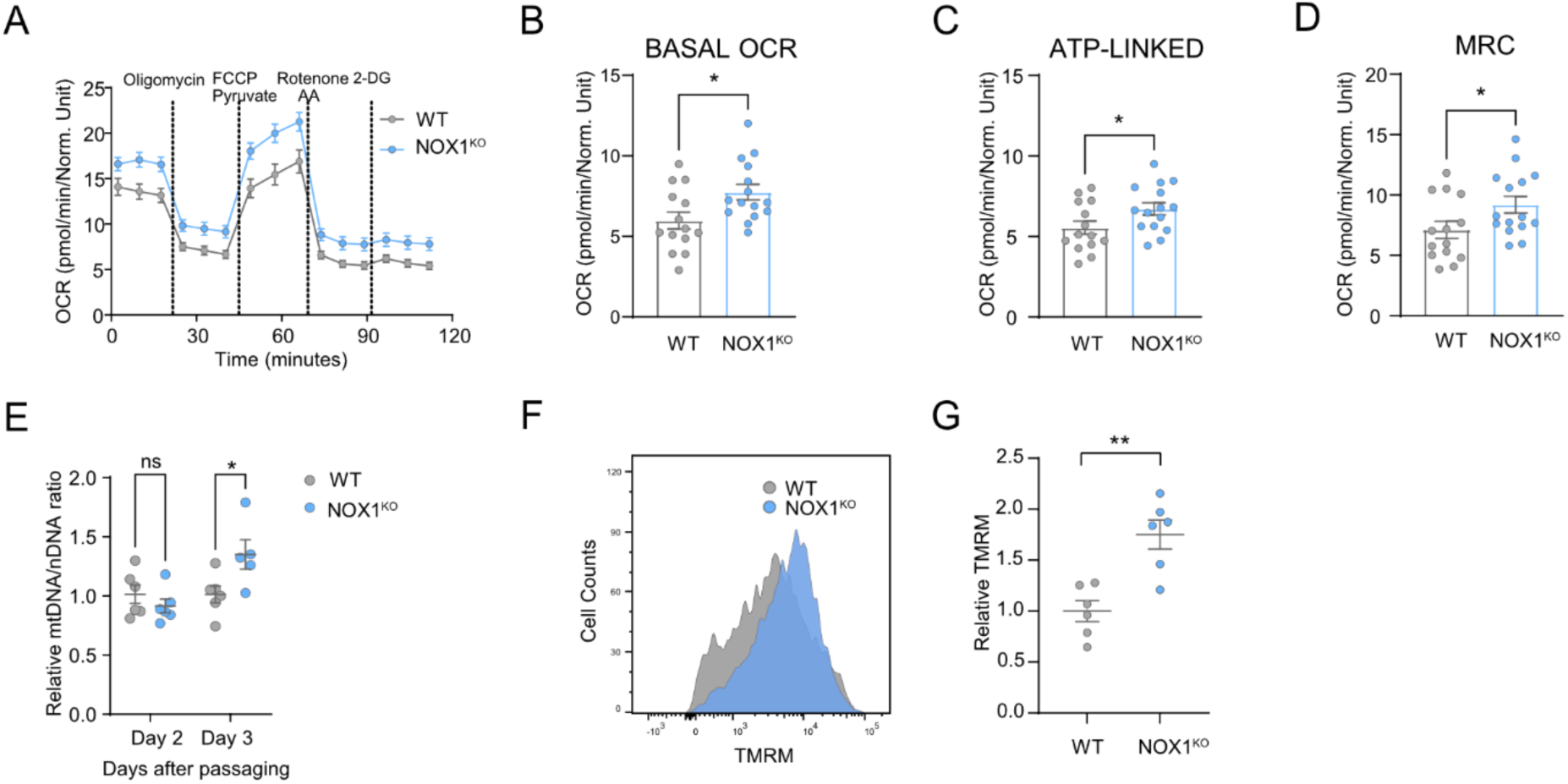
NOX1 deficiency enhances OXPHOS activity in cycling IECs. (A-D) Kinetic line graph (A) of oxygen consumption rate (OCR) in 2D epithelial monolayers from WT and NOX1^KO^ littermates measured by a Seahorse Analyzer. (B) is basal OCR, (C) is ATP-linked respiration and (D) is MRC bar graphs. Each dot indicates one biological replicate (n=3 mice). (I) qRT-PCR for mitochondrial ND1 and 16S rRNA gene expression in colonoids from WT and NOX1^KO^ littermates collected on indicated day after switched to non-Wnt ERN medium. Values were normalized to *Rpl13a*. Each dot indicates one biological replicate (n=3 mice). (F and G) Representative tetramethylrhodamine methyl ester (TMRM) FACS plots (F) and quantification of relative geometric mean fluorescence intensity of TMRM (G) in single cells of colonoids from WT and NOX1^KO^ littermates cultured in Wnt-rich WERN medium. Values were normalized to WT colonoids. Each dot indicates one biological replicate (n=3 mice). Experiments were performed no less than 3 times. Data (A), (B), (C), (D), (E) and (G) are presented as mean ± SEM. Statistical significance was determined using two-tailed unpaired Student’s t tests in (B), (C), (D) and (G), Two-way ANOVA in (E); ns, not significant; *p < 0.05 and **p < 0.01.

**Figure S7.**
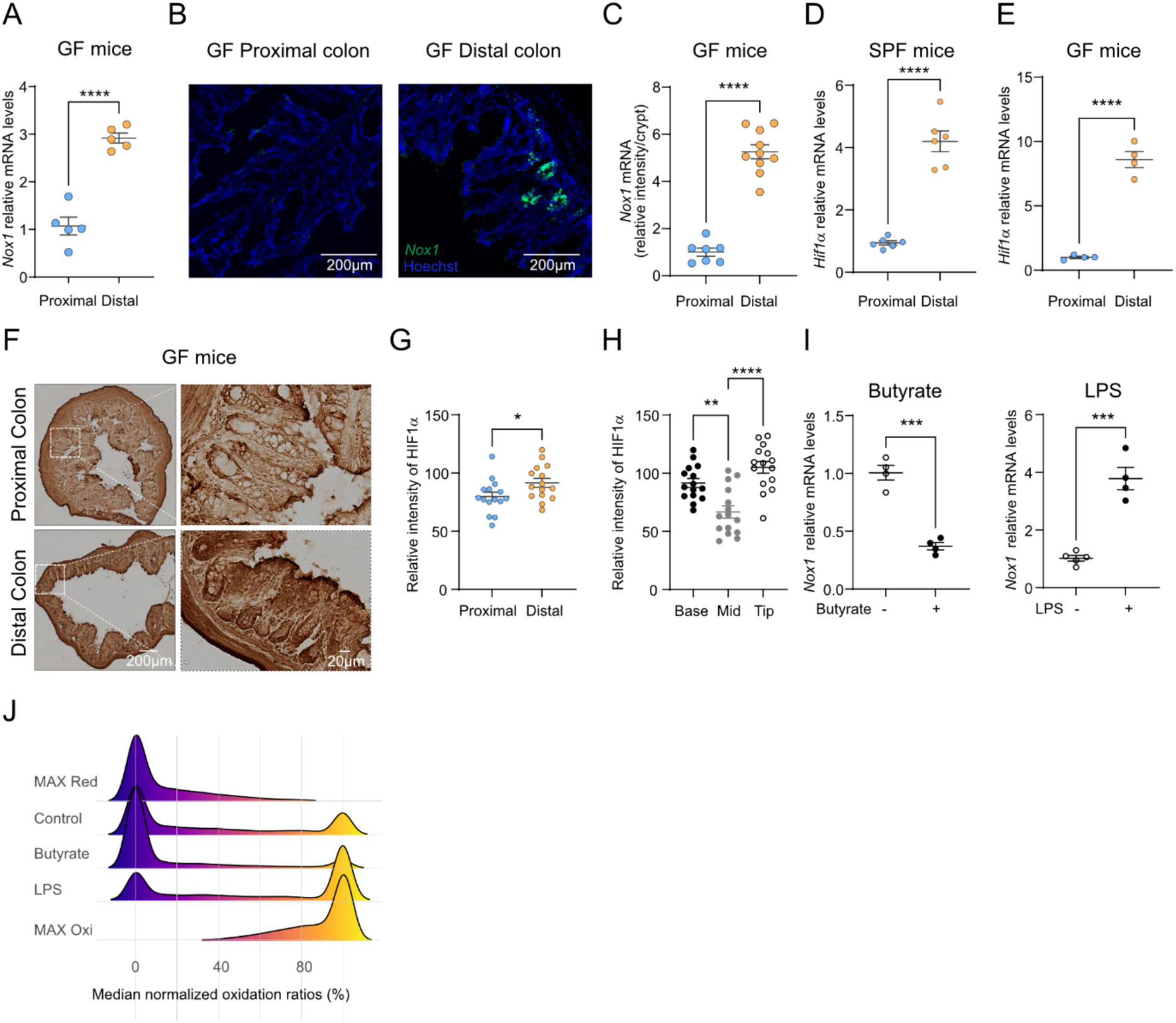
Niche-dependent *Nox1* and HIF1α expression patterns are not microbe dependent. (A) qRT-PCR for *Nox1* expression in proximal colons and distal colons from GF mice. Values were normalized to *Rpl13a*. Each dot indicates one mouse. (B and C) RNAScope images (B) and quantification of relative *Nox1* mRNA intensity (C) of vertical sections of freshly frozen proximal colons and distal colons from GF mice, probed for *Nox1* mRNA in green. Each dot indicates one ROI (2-3 ROI/mouse, n=3 mice). (D) qRT-PCR for *Hif1α* expression in proximal colons and distal colons from SPF mice. Values were normalized to *Rpl13a*. Each dot indicates one mouse. (E) qRT-PCR for *Hif1α* expression in proximal colons and distal colons from GF mice. Values were normalized to *Rpl13a*. Each dot indicates one mouse. (F-H) DAB analysis (F) and quantifications (G and H) of HIF1α accumulation in proximal and distal colon FFPE samples. Each dot indicates one ROI (5 ROI/mouse, n=3 mice). (I) qRT-PCR for *Nox1* expression in colonoids from distal colons of R26-RoGFP2-Orp1^Villin^ mice cultured in Wnt-rich WERN medium, treated with or without 1 mM butyrate, 10 μg/ml LPS treatment separately. Each dot indicates one biological replicate (n=3 mice). (J) Oxidation of roGFP in response to treatment indicated in (I). Maximum reduction (purple) and oxidation (yellow) of roGFP following additions of 2 mM DTT or 10 μM diamide respectively are shown as a reference. Experiments were performed no less than 3 times. Scale bars, as indicated. Data (A), (C), (D), (E), (G), (H) and (I) are presented as mean ± SEM. Statistical significance was determined using two-tailed unpaired Student’s t tests in (A), (C), (D), (E), (G) and (I), One-way ANOVA in (H); *p < 0.05, **p < 0.01, ***p < 0.001 and ****p < 0.0001.

**Figure S8.**
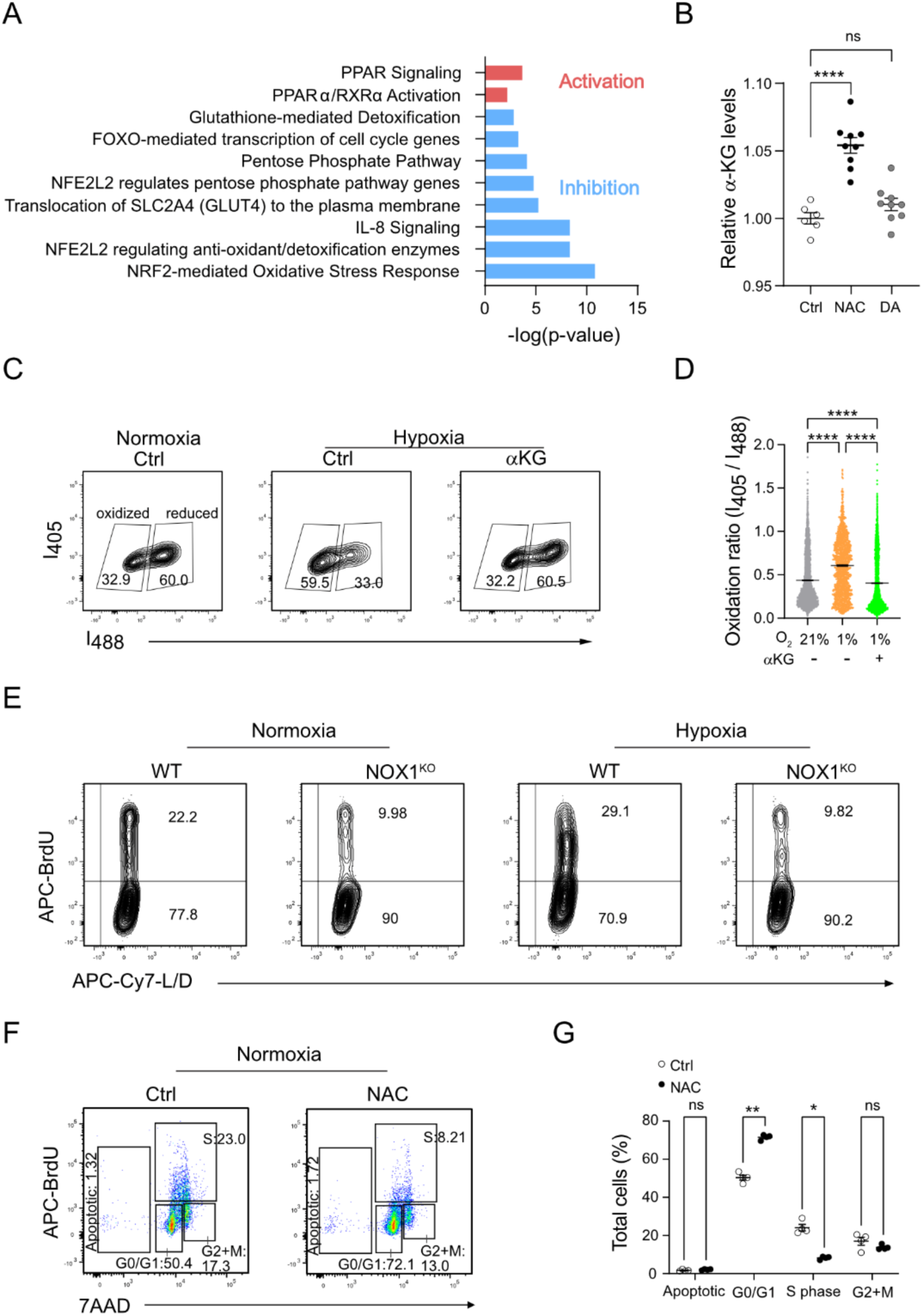
Redox status regulates metabolism and hypoxia response during cell cycle progression in cycling IECs. (A) Ingenuity Pathway Analysis of Figure 7A. (B) α-Ketoglutarate levels assay of colonoids from distal colons of WT treated with treated with or without 5 mM NAC or 10 μM diamide. Each dot indicates one biological replicate (n=3 mice). (C and D) Flow cytometry fluorescence measurements (C) and quantification (D) of 405/488-nm ratios in distal colon-derived colonoids of R26-RoGFP2-Orp1^Villin^ mice under normoxia (21% O_2_), or hypoxia (1% O_2_), treated with or without 3 mM α-Ketoglutarate. Each dot indicates one cell (n=3 mice). (E) Representative flow plots of BrdU incorporation in distal colon-derived colonoids of WT and NOX1^KO^ littermates under normoxia (21% O_2_), or hypoxia (1% O_2_) for 2 h. See also Figure 7G. (F and G) Representative flow plots (F) and quantification (G) of cell cycle analysis in WT colonoids under normoxia (21% O_2_), treated with or without 5 mM NAC. Each dot indicates one biological replicate. Experiments were performed no less than 3 times. Data (B), (D) and (G) are presented as mean ± SEM. Statistical significance was determined using One-way ANOVA in (B) and (D), Two-way ANOVA in (G); ns, not significant; *p < 0.05, **p < 0.01 and ****p < 0.0001. (F)

## STAR METHODS

### Key Sources Tables

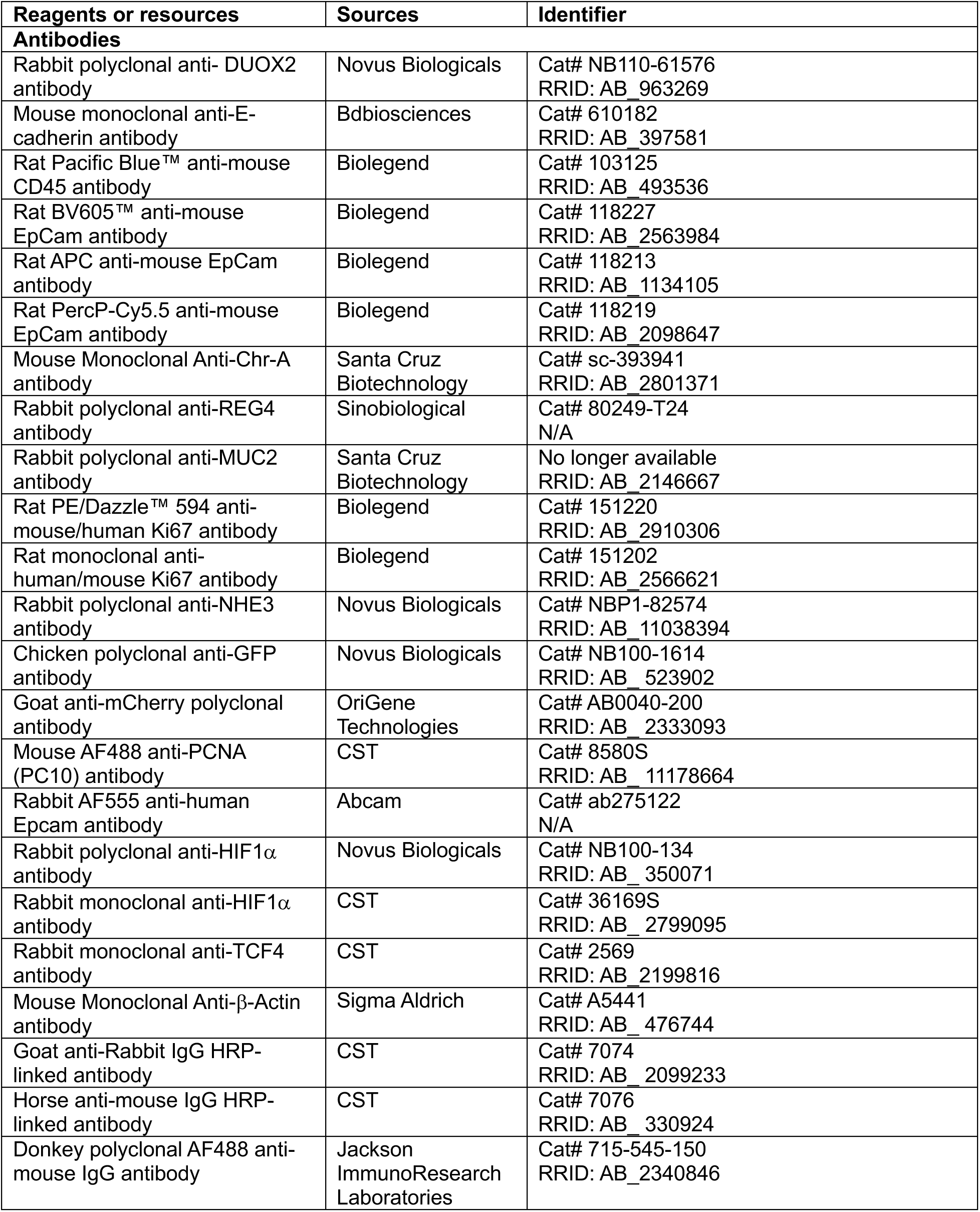
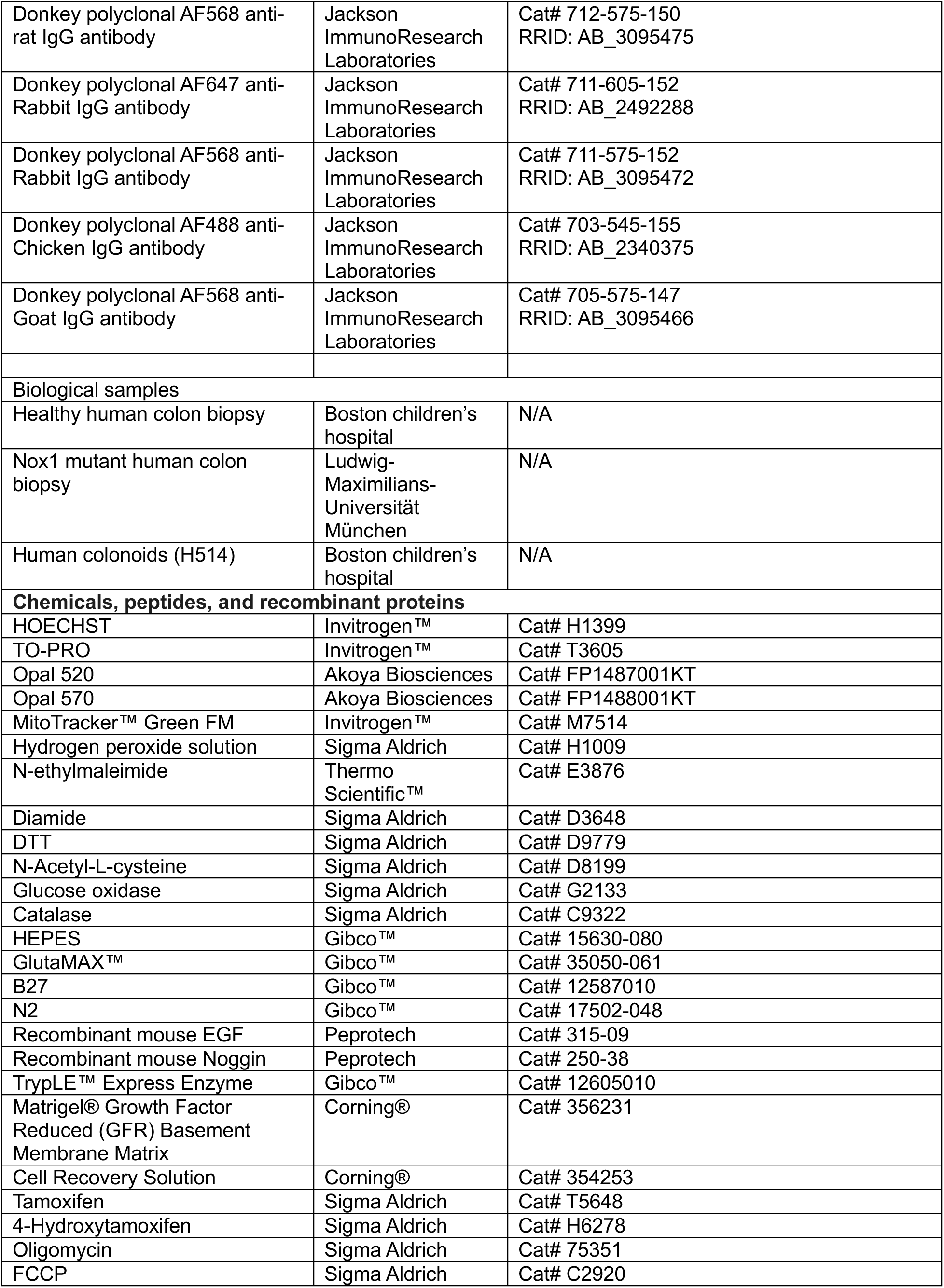
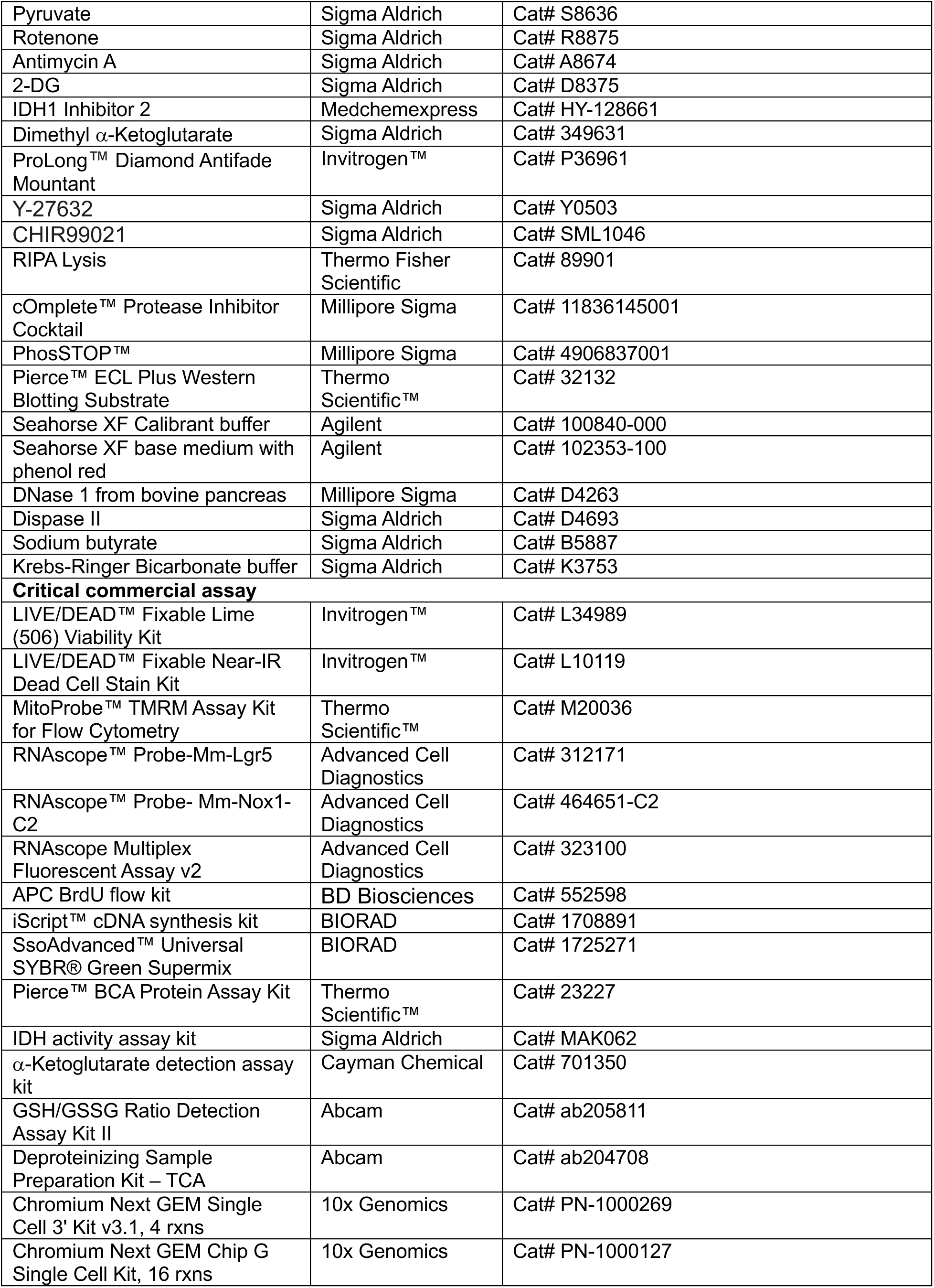
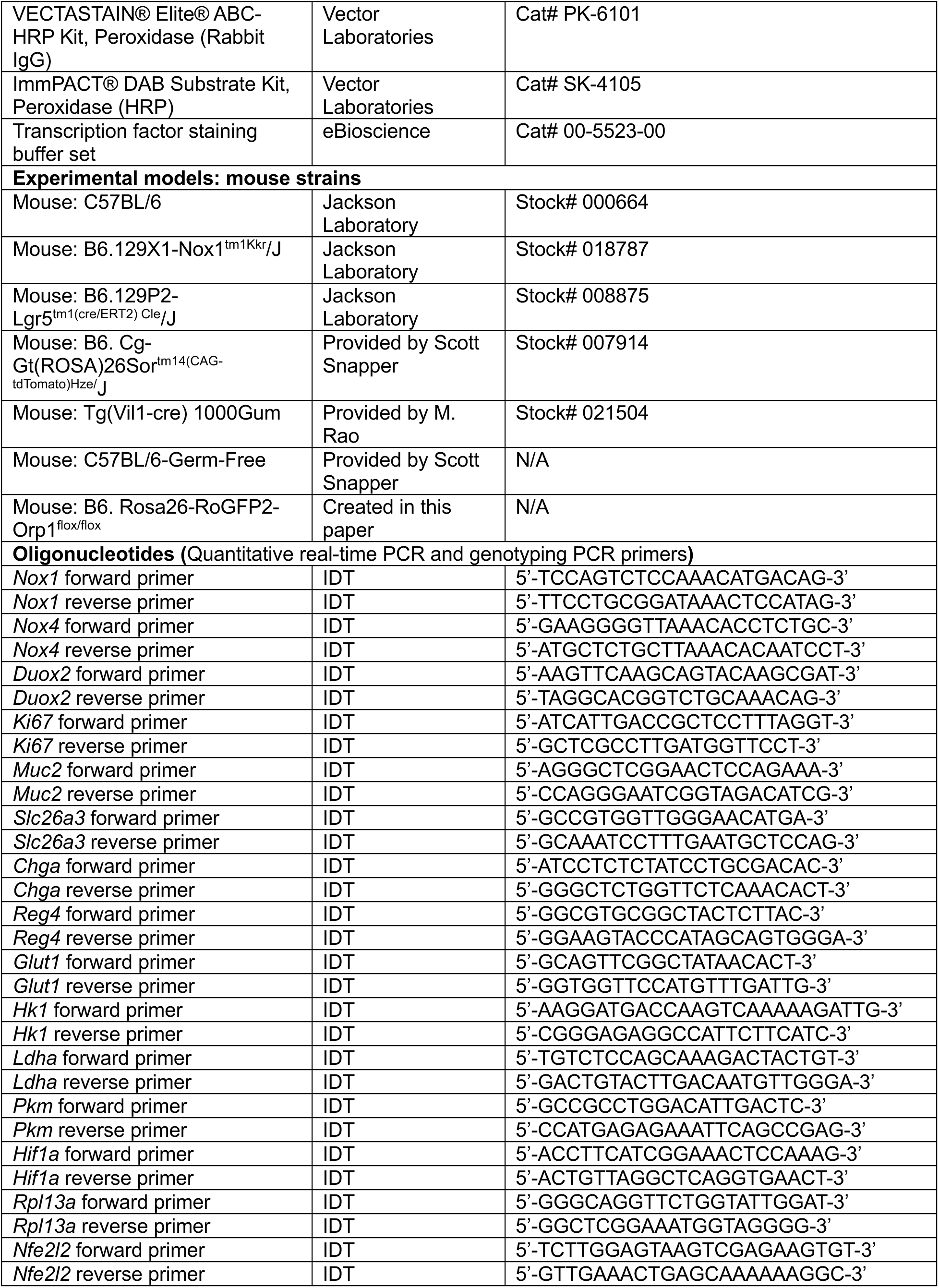
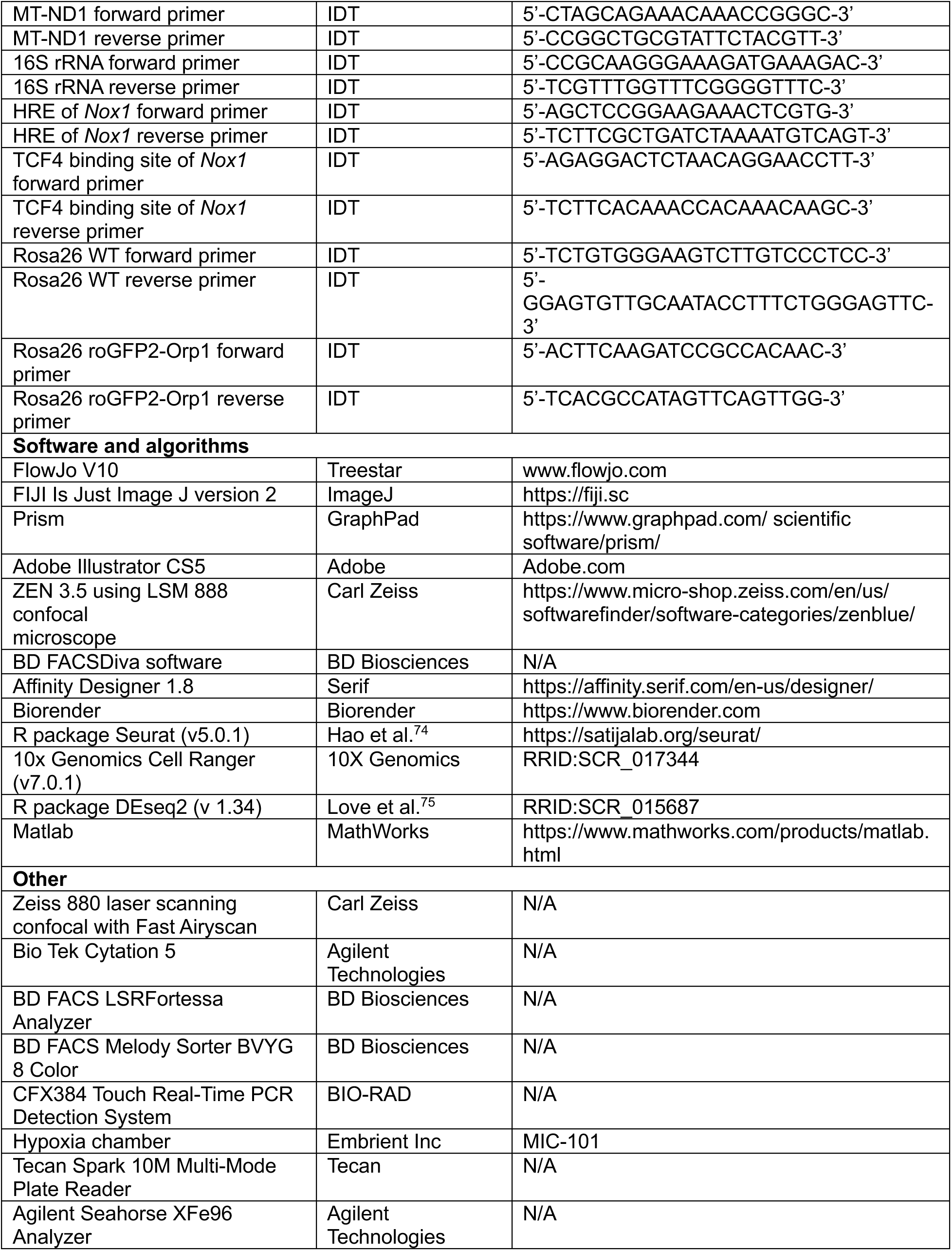

## EXPERIMENTAL MODEL AND SUBJECT DETAILS

### Animals

C57BL/6, B6.129X1-Nox1^tm1Kkr^/J (NOX1KO) and B6.129P2-Lgr5^tm1(cre/ERT2) Cle^/J (Lgr5^creERT2/+^) mice were purchased from The Jackson Laboratory. B6.Cg-Gt(ROSA)26Sor^tm14(CAG-tdTomato)Hze^/J (Rosa26^tdtomato^) strain and germ free mice were gifted from Snapper Scott Lab. GF mice were bred and maintained in vinyl isolators in the Animal Resources at Children’s Hospital (ARCH). Stool samples from GF were routinely collected and tested for sterility and exclusion of other bacterial contamination using aerobic and anaerobic conditions culture. Tg(Vil1-cre) 1000Gumstrain was gifted from Meenakshi Rao Lab. To explore the role of NOX1 in intestinal stem cells, we crossed NOX1^KO^ mouse strain and Lgr5^creERT2/+^ to generate NOX1^KO^; Lgr5^creERT2/+^ mouse strain. To generate genetic lineage tracing strain, we bred Lgr5^creERT2/+^ and Rosa26^tdtomato^ to generate Lgr5^creERT2/+^; Rosa26^tdtomato/+^mouse strain. To explore the role of NOX1 in cell fate decision, we bred NOX1^KO^ mouse strain with Lgr5^creERT2/+^; Rosa26^tdtomato/+^ to generate NOX1^KO^; Lgr5 ^tdtomato^ mice.

To generate intracellular hydrogen peroxide reporter mice R26-RoGFP2-Orp1^flox/flox^ using the CRISPR-Cas9 method, we used the cloning-free CRISPR/Cas system.^76^ pR26roGFP2Orp1 targeting vector was generated by assembling Orp1-roGFP2 to the AsiSI/MluI restriction sites in Rosa26 targeting vectors pR26 CAG AsiSI/MluI.^77^ 0.61 pmol each of crRNA and tracrRNA was conjugated and incubated with Cas9 protein (30 ng/μl) to prepare ribonucleoprotein complex (RNP) and donor DNA (10 ng/μl) was mixed with RNP for microinjection cocktail.^2^ Microinjection cocktail was injected into 0.5 dpc embryos harvested after mating C57Bl6-Hsd (Envigo) females. Post-injection embryos were reimplanted into CD1 (Envigo) pseudo-pregnant foster females and allowed to term. Tail snip biopsies were collected from pups at P7 for genotyping. To generate intestinal epithelial cell specific intracellular hydrogen peroxide reporter mice, we crossed R26-RoGFP2-Orp1flox^/flox^ mice with Tg(Vil1-cre) 1000Gummice to get R26-RoGFP2-Orp1^Villin^ mice.

Mice were bred and housed in a specific pathogen-free animal facility of Animal Resources at Children’s Hospital (ARCH). Age-matched 6- to 12-week-old littermate male and female mice were used for experiments. All animal experiments were approved by the Children’s Institutional Animal Care and Use Committee. Mice were maintained in 12 hours light/dark cycle, food and water was provided *ad libitum*.

### Human intestinal tissue samples

Healthy human sigmoid biopsy tissue was obtained from a 5-year-old male patient during diagnostic colonoscopy as part of routine clinical care under approved Boston Children’s Hospital Institutional Review Board Protocol #P00046566. *Nox1* mutant p.N122H human colonic biopsy tissue was obtained from a 8-year-old male patient as a part of a study approved by the institutional review board of the Ludwig-Maximilians-Universität München, and samples were collected upon informed consent. Biopsy samples were fixed in 10% Neutral Buffered Formalin, embedded in paraffin, and cut into 5-7 μm sections for staining.

### Primary cell cultures

Murine colonic organoids were generated from mouse colonic crypts. Murine colonic organoids were maintained at 37°C in a humidified atmosphere containing 5% CO_2_. Colonoids were plated in a 50 ml drop of Matrigel and overlaid with 500 ml of WENR medium, containing basal crypt media (Advanced DMEM/F12, penicillin/streptomycin, 10 mM HEPES, 2 mM Glutamine), supplemented with 1× B27 (Gibco, 17504-044), 1× N2 (Gibco,17502-048), 50 ng/ml rmEGF (Peprotech, 315-09), 50% L-WRN-CM (conditioned medium) (v/v), 10 mM Rock inhibitor Y-27632 (Sigma, Y0503), 250 nM CHIR99021 (Sigma, SML1046) with a final FBS concentration of 10%. Human colonoids H514 were generated from a non-diseased pediatric colonic biopsy. Human colonoids were maintained at 37°C in a humidified atmosphere containing 5% CO_2_. Colonoids were plated in a 50 ml drop of Matrigel and overlaid with 500 ml of WENR medium, containing basal crypt media (Advanced DMEM/F12, penicillin/streptomycin, 10 mM HEPES, 2 mM Glutamine), supplemented with 1× B27 (Gibco, 17504-044), 1× N2 (Gibco,17502-048), 50 ng/ml rhEGF (Peprotech, AF-100-15), 50% L-WRN-CM (conditioned medium) (v/v), 10 mM Nicotinamide (Sigma, N0636), 500 nM A83-01 (Sigma, SML0788), 500 nM SB202190 (Sigma, S7067), 500 mM N-Acetyl-Cysteine (Sigma, A8199), 50 nM Gastrin (Sigma, G9145), 100 nM Prostaglandin E2 (Sigma, P5640), 10 mM Rock inhibitor Y-27632 (Sigma, Y0503), 250 nM CHIR99021 (Sigma, SML1046) with a final FBS concentration of 10%.

## METHOD DETAILS

### 3D organoids primary cell culture and treatment

Colonic organoids were generated from isolated crypts collected from distal colon of mice listed above as previously described.^78^ Approximately 500 crypts were plated in a 50 μl drop of Matrigel in 24-well plates and overlaid with 500 μl of WENR medium, containing basal crypt media (Advanced DMEM/F12, 100 U/ml penicillin/streptomycin, 10 mM HEPES, 2 mM Glutamine), supplemented with 1× B27 (Gibco, 17504-044), 1× N2 (Gibco,17502-048), 50 ng/ml rmEGF (Peprotech, 315-09), 50% L-WRN-CM (conditioned medium) (v/v), 10 μM Rock inhibitor Y-27632 (Sigma, Y0503), 250 nM CHIR99021 (Sigma, SML1046) with a final FBS concentration of 10%. Media was changed every 2 days and colonoids were split every 4 days. Cultures were passaged three times prior to experiments. For the measurement of spheroids surface area and the calculation of organoids formation number, we seeded 1000 cells/small cell clumps per well, 24 hrs after seeding, colonoids were treated with or without 1 mM NAC or 10 μM diamide and then we took images with a BioTek Cytation5 automated microscope at 2.5X object on day 3 or 5 after plating. The percentage of stem-like spheroids versus budded colonoids, were manually counted based on brightfield images (BioTek Cytation5 at 2.5X) after 3 days following plating in and treated with or without 200U catalase (Sigma C9322) or 2000U glucose oxidase/20mM glucose (G2133). To induce colonoids differentiation, 24 h after seeding, primary epithelial cells were exposed to ENR medium containing basal crypt media (Advanced DMEM/F12, penicillin/streptomycin, 10 mM HEPES, 2 mM Glutamine), supplemented with 1× B27 (Gibco, 17504-044), 1X N2 (Gibco,17502-048), 50 ng/ml rmEGF (Peprotech, 315-09), 100 ng/ml Noggin (Peprotech 250-38), 10% R-spondin1-CM (conditioned medium) (v/v) with a final FBS concentration of 10%. For the stimulation with different niche factors, 24 hrs after seeding, primary epithelial cells were exposed to either ENR medium, or in WENR medium with or without butyrate (1 mM), LPS (10 μg/ml), antioxidant DTT (2 mM) or oxidant diamide (10 μM) treatment or were cultured either under 21% oxygen (normoxia) or 1% oxygen (hypoxia) conditions in WENR or ENR medium.

### RNA isolation and real-time QPCR analysis

Lyse and homogenize snap frozen tissues or colonoids in Matrigel in TRIzol™ Reagent. Then isolate RNA according to manufacturer’s instructions. Equivalent amounts of purified RNA was converted to cDNA using iScript cDNA synthesis kit (Bio-rad) and analyzed for gene expression on a CFX384 real time cycler (Bio-rad) using iTaq Universal SYBR Green Supermix (Bio-rad) with Qpcr primers from Primer Bank for the following genes: *Nox1, Nox4, Duox2, Ki67, Muc2, Slc26a3, Chga, Reg4, Glut1, Hk1, Ldhα, Pkm, Hif1α,* which are listed in the Key Sources Table. Data were normalized to *Rpl13a*, and results were shown as fold induction relative to expression levels in WT groups or control groups.

### RNAScope

For mouse *Nox1* and *Lgr5* RNAscope staining, proximal colons and distal colons were collected, embedded in Tissue-Tek OCT compound and snap-frozen in dry ice before freezing at −80 °C. Embedded tissues were warmed to −20 °C, cryosectioned into 14mm slices and then were fixed with 4% PFA at 4 °C for 15 min and stained for *Nox1* and *Lgr5* using the RNAscope Multiplex Fluorescent Assay v2 (Advanced Cell Diagnostics) following the manufacturer’s instructions. Entire sections were imaged using a confocal microscopy at 20X and images were analyzed in FIJI.

### Tissues preparation and images acquisition from intracellular hydrogen peroxide reporter mice (Colon tissues/colonoids)

For freshly fixed frozen samples, proximal colons and distal colons were embedded in Tissue-Tek OCT compound and snap-frozen in dry ice before storage at −80 °C. Embedded tissues were warmed to −20 °C, cryosectioned into 14 mm slices with a cryotome (Leica CM3050) and mounted onto Superfrost Plus slides (Fisher Scientific GmbH). For sections further processed with chemicals, the chemicals were applied immediately after mounting. To induce maximal probe oxidation or reduction, sections were incubated for 10 min on ice with 1 mM DA or 20 mM DTT, respectively, before NEM treatment. To preserve the endogenous redox state, sections were immediately incubated with 50 mM NEM (Sigma-Aldrich, E3876) for 10 min on ice followed by fixation in 4% PFA in PBS containing 1 mM TO-PRO-3 (using 0.1% dimethyl sulfoxide as a co-solvent) (Invitrogen, T3605) for 15 min at ambient temperature. PFA was washed out twice with ice-cold PBS for 5 min. Sections were then mounted in ProLong Dimond Antifade Mountant (Thermo Fisher Scientific) and kept at 4 °C until use.^21^

Guidelines for the use of roGFP2-based redox probes, including microscopy settings and image analysis, have been provided previously.^21,79,80^ Adapted from that, images were acquired using Zeiss LSM 880 using 40X 1.3 NA objective. For each animal, at least three images were taken from one or more sections. Three channels were imaged (2048×2048 pixels at 103 nm resolution). Nuclei stained with TO-PRO 3 was imaged in the far-red channel with the 633 nm excitation laser. For measuring the redox changes, two excitation lasers (405 nm and 488 nm) were used to sequentially image an emission channel (500-540 nm). Images were analyzed using custom written MATLAB code to generate the redox ratio metric images (I_405_/I_488_). Each channel was first eroded using a structural element of 5 pixels. An intensity threshold of 7 was used to denoise the image and the ratio of the channels was calculated. Any ratio greater than 20 was set to 0 to prevent low signal regions. For each image, the median of the non-zero ratio values is calculated. Final data were plotted using custom R programs. For live imaging, harvested tissues or colonoids in Matrigel were mounted in a live imaging chamber in Krebs-Hepes buffer (pH7.2) and imaged on a Zeiss LSM 880 microscope using a 20x 0.8NA objective.

### Colon tissue Immunofluorescence microscopy

For FFPE samples, tissues were fixed with 4% paraformaldehyde diluted in PBS at 4 °C overnight. Tissues were processed, paraffin-embedded, and cut into 5 μm sections at BIDMC histology core. 5 μm tissue cross-sections were heated at 60 °C for 10 min prior to deparaffination in xylene for 10 min twice, and then rehydrated with decreasing concentrations of ethanol (100, 95, 70, 50 and 0%) for 10 min each at room temperature. Antigen retrieval was performed by pressure cooker in antigen retrieval solution (10 mM citric buffer, pH 6.0) for 15 min on high. Samples were washed in PBS and blocked with 10% donkey serum in PBS for 1 h at 4 °C in a humidified chamber. Then samples were incubated with the indicated primary antibodies diluted in blocking buffer at 4 °C overnight. Sections were washed twice and then stained with fluorescent secondary antibodies diluted in blocking buffer for 2 hrs at room temperature. Finally, the sections were washed three times and incubated with PBS containing Hoechst at a 1:1,000 dilution, and then mounted with ProLong Dimond Antifade Mountant (Thermo Fisher Scientific).

For cryosection samples, colons were fixed with 4% paraformaldehyde diluted in PBS for 1-2 hours at 4 °C, washed with PBS three times, incubated with 30% sucrose at 4 °C overnight, and then embedded in OCT (Tissue Tek), snap-frozen in dry ice before storage at −80 °C. Embedded tissues were warmed to −20 °C, cryosectioned into 14 mm slices with a cryotome (Leica CM3050) and mounted onto Superfrost Plus slides (Fisher Scientific GmbH). Samples were then blocked with 10% donkey serum in PBS for 1 h at 4 °C in a humidified chamber and were then incubated with the indicated primary antibodies diluted in blocking buffer at 4°C overnight. Sections were washed twice and then stained with fluorescent secondary antibodies diluted in blocking buffer for 2 hrs at room temperature. Finally, the sections were washed three times and incubated with PBS containing Hoechst at a 1:1,000 dilution, and then mounted with ProLong Dimond Antifade Mountant (Thermo Fisher Scientific). For each animal, at least two-three images were taken from one or more sections with a 20X objective on a Zeiss LSM 880 inverted confocal microscope, some images were under automated tile scanning with 10% overlap between individual images.

### Organoids tissue Immunofluorescence microscopy

After recovered from Matrigel, colonoids were incubated in 4% paraformaldehyde diluted in PBS with gentle rocking for 45 min at 4°C. Let the colonoids settle by gravity and wash the colonoids once in immunofluorescence (IF) buffer, PBS containing 0.1% w/v BSA, 0.2% v/v TritonX-100, (0.05% v/v) TWEEN^®^ 20. We then resuspended the colonoids in 1 ml of PBS and transfer them to 1.5 ml Eppendorf tubes. Aspirate PBS and add 1 ml of citrate buffer (pH 6.0) to the colonoids and incubate the tubes in the heating block of 98 °C for 20 minutes. Then aspirate the citrate buffer and add 1 ml of permeabilization/blocking solution, PBS containing 5% v/v serum, 1% v/v TritonX-100, and incubate at room temperature with agitation for 2 hours. Aspirate the permeabilization/blocking solution and wash 3 times in IF Buffer. Add 0.5 ml of the primary antibodies diluted in IF Buffer and incubate at room temperature overnight with gentle agitation. Aspirate the primary antibody solution and wash 3 times in IF Buffer. Add 0.5-1 ml of the secondary antibodies diluted in IF Buffer supplemented with 10% donkey serum and incubate at room temperature for 2 hours with gentle agitation. Add Hoechst directly to the secondary solution/colonoids to a final concentration of 2-4 μg/ml and incubate at room temperature with gentle agitation for 15-20 min in the dark. Aspirate the staining solution and wash the colonoids once in water and once in PBS and then resuspend the organoid pellet in 50 μl ProLong™ Gold Antifade Mountant. Dispense the organoid/ mountant suspension onto the center of a glass microscope slide within image spacers and cover with a coverslip. Images were taken with a 20X objective on a Zeiss LSM 880 inverted confocal microscope. This was adjusted from protocol of Stem Cell Technologies.

### Flow cytometry

To determine the intracellular hydrogen peroxide levels in colonoids, we digested colonoids with TripLE to get single cell suspensions. After stained with live/dead dye, cells were immediately treated with 50 mM NEM to clamp the intracellular redox status. roGFP 405/488 ratio analysis by FACS was performed in a Fortessa (BD Biosciences) equipped with a 405 nm and 488 nm laser lines. The cells were excited with 405 nm, and 488 nm lasers and fluorescence emission were collected at 525/50 nm or 530/30 nm at each laser line in ratio mode. The oxidation degree of roGFP was calculated adapted from Avia et al and Schwarzländer et al.^81,82^ Flow data was analyzed using custom written R code to calculate median normalized oxidation ratio. The 405 to 488 ratio was calculated for each cell and the outliers removed using interquartile range as a measure. All the ratios were normalized to their respective median values and scaled using the median value of the DTT treatment (normalized as 0%) and the median value of the DA treatment (normalized as 100%).

To explore the role of NOX1 in ISCs regeneration, we did BrdU incorporation in both mice and *in vitro* colonoids. For mice experiments, we injected mice IP with 100 μl (1 mg) of BrdU solution (BD Pharmingen). 2 hours later, we sacrificed the mice and isolated colonic crypts with 5 mM EDTA, then digested them with dispase II and DNaseI to get single cell suspensions. For organoids culture, we incubated cells with BrdU at a final concentration of 10 μM in cell culture medium for 1 h. After recovered from Matrigel, colonoids were digested with TripLE into single cell suspensions. Live/dead dye, mouse anti-CD45 Pacific Blue antibody, mouse anti-EpCam BV605 antibody, anti-BrdU APC antibody was used. Then cells were stained following the manufacturer’s instructions. For cell cycle analysis, cells were stained with 7AAD that binds to total DNA after anti-BrdU staining. Data were acquired on a BD LSRFortessa or a BD Symphony A5 SE spectral flow cytometer and analyzed in Flowjo v10.

For genetic lineage tracing *in vivo*, mice were injected with tamoxifen 20 mg/kg and sacrificed 48 hrs after induction. We then isolated colonic crypts with 5mM EDTA, digested them with dispase II and DNaseI to get single cell suspension. Then dissociated cells were stained with live/dead dye, mouse anti-CD45 Pacific Blue, mouse anti-EpCam BV605 and analyzed by BD LSRFortessa analyzer.

For intestinal stem cell isolation, colonic crypts were isolated with 5mM EDTA from proximal and distal colons of WT; Lgr5^creERT2+^ and NOX1^KO^; Lgr5^creERT2+^ mice. Then crypt pellets were digested with dispase II and DNase I to get single cell suspension. Then dissociated cells were stained with live/dead dye, mouse anti-CD45 Pacific Blue, mouse anti-EpCam PercP-CY5.5, and sorted with BD Melody sorter. Cells were sorted depending on EGFP intensity. Primary Organoids formation analysis was performed with sorted intestinal stem cells.

For intestinal epithelial cells (IECs) isolation for single cell sequencing, we isolated colonic crypts from distal colons and proximal colons of WT and NOX1^KO^ littermates with 5mM EDTA, digested them with dispase II and DNaseI to get single cell suspension. Then the dissociated cells were stained with live/dead dye, mouse anti-CD45 Pacific Blue, mouse anti-EpCam PercP-CY5.5, and IECs were sorted with BD Melody sorter.

### Fasting-refeeding models

During the fasting period, 8-10-week-old mice were placed on a stainless mesh floor with no food to avoid coprophagia, and with ad libitum drinking water. Mice were fasted for 36 hrs and then refed with normal chow for 24 hrs. Mice were monitored every 12 hrs for body weight change.^23^ Colons were harvested for flow cytometry analysis or tissue H&E staining, other immunohistochemical staining.

### 10X scRNA-seq of mouse colon

Colonic epithelial cells isolated from WT and NOX1^KO^ littermates by BD Melody sorter from proximal colon or distal colon regions were pooled separately across 2 mice at equal ratios and were resuspended in 1% BSA in PBS and counted with trypan blue staining for cell viability and number. Approximately 30,000 cells per sample were loaded onto separate lanes of the Chromium X Controller. Gene expression libraries were prepared according to manufacturer instructions using the 10X Genomics Single Cell 3’ Reagent Kit (v3.1 Chemistry Dual Index). Sequencing was performed on a NovaSeq 6000 through the Broad Clinical Labs using a NovaSeq S2 100 cycle kit to target 20,000 read pairs per cell.

### 10X scRNA-seq data processing and downstream analysis

Sequencing files were processed using CellRanger (v7.2.0). Analysis was conducted using Seurat (v5.1.0)^74^ keeping cells containing at least 100 unique genes and features expressed in at least 3 cells. A coarse quality filter was applied to remove cells with greater than 8000 unique genes and 50% of reads mapping to mitochondrial genes. The expression and metadata were converted into Seurat objects, one for proximal and other for distal colon. Gene counts were scaled and normalized using log normalization and the 3000 most variable features selected after variance stabilization were then used in dimensionality reduction by PCA. 30 PCs were used for UMAP dimensional reduction and a granularity of 0.2 and 0.5. The top 10 markers for each cluster were determined and the clusters manually annotated.

Trajectory analysis was performed using Monocle 3.0 with a cluster resolution of 5e-4 and the cluster corresponding to the Stem Cell population was used as the root cell. The low-quality, enteroendocrine and tuft cell clusters were removed from this analysis. 3.5 was used as a cutoff to delineate high and low Transit Amplifying cells. A manually curated list of cell cycle genes was used to generate relative expression under different conditions for the TA cell.^83^ Data can be accessed at https://www.ncbi.nlm.nih.gov/geo/query/acc.cgi?acc=GSE289397.

### Seahorse cellular metabolic analysis

One day prior to assay, transform 3D organoids to 2D monolayer. Colonoids were extracted from Matrigel with organoid harvesting solution (R&D system, 3700-100-01), and then digested with undiluted TripLE (Life Technologies, 1265010). Colonoids fragments were suspended in WENR medium and plated on collagen-coated (Sigma, C5533) Seahorse 96-well microplate plate. Meanwhile, hydrate cartridge and warm Seahorse XF Calibrant. After 24 hours, cells were washed twice and incubated in the Seahorse Assay Medium supplemented with 12 mM glucose and 2 mM glutamine at 37 °C for 45 min. Oxygen consumption rate (OCR) and extracellular acidification rate (ECAR) were measured with a Seahorse XFe96 Extracellular Flux Analyzer. The OCR and ECAR were measured under basal conditions and after injection of OM (2 μM), FCCP (2 μM) plus pyruvate (5 mM), rotenone (1 μM) plus antimycin A (1 μM) (Rot.+AA), and 2-DG (50 mM). Metabolic parameters were calculated as follows: Basal OCR = OCR_before OM_ - OCR_after Rot+AA_, ATP-linked respiration = Basal OCR - OCR_after OM_, Maximal Respiratory Capacity (MRC) =OCR_after FCCP+Pyruvate_ - OCR_after Rot+AA_, Basal ECAR = ECAR_before OM_ - ECAR_after 2-DG_, Maximal Glycolytic Capacity (MGC) = ECAR_afterOM_ - ECAR_after 2-DG_.

### Mitochondrial membrane potential measurement by flow cytometry

Colonoids were treated with TMRM (20 nM) and MitoTracker Green (20 nM) for 30 min at 37°C. After recovered from Matrigel, colonoids were digested with TripLE into single cell suspensions. Then dissociated cells were stained with live/dead dye and then analyzed by BD LSRFortessa flow cytometer with 561-nm excitation.

### Bulk RNA-seq of colonoids

Lyse colonoids recovered from Matrigel in TRIzol™ Reagent. Then isolate RNA according to manufacturer’s instructions. The quality of RNA samples was determined using an Agilent 5400 Bioanalyzer, and all samples for sequencing had RNA integrity (RIN) numbers >8. cDNA library construction using the NEBNext^®^ Ultra^™^ II RNA Library Prep Kit for Illumina^®^ was performed by the Novogene Corporation Inc. (Sacramento, CA), and cDNA libraries were sequenced on an Illumina HiSeq 2000 or Illumina Novaseq 6000 instruments.

### Bulk RNA-seq data analysis

Quality control for mRNA-seq data (FASTQ files) was conducted using FastQC (v0.11.9) followed by alignment to the mouse genome (GRCm38.p6) using the STAR aligner (v2.7.8a) in 2-pass mode with default parameters and version 25 of the mouse GENCODE primary assembly annotation (GENCODEvM25). Aligned BAM files were further analyzed using and RSeQC (v4.0.0). The alignments and quality control steps were run using the Snakemake workflow management system (v6.6.1). Reads belonging to individual genes annotated by GENCODEvM25 were counted using htseq-count (v0.13.5). Differential expression analysis was run using the default workflow provided by the DESeq2 R package (v1.34.0). Volcano plot was charted using the Enhanced Volcano program.^84^ A set of 128 downregulated genes that were on average at least 50% higher in control relative to KO was used to determine pathways altered as a consequence of knockdown.

Data can be accessed at https://www.ncbi.nlm.nih.gov/geo/query/acc.cgi?acc=GSE289209

### Oxidative Isotope-Coded Affinity Tags (OxICAT) sample preparation and LC-MS/MS analysis

OxICAT sample preparations were adapted from Bechtel, T. *et al*^85^ and Topf, U. *et al*.^86^ 100 μg of cell lysates of colonoids recovered from Matrigel was precipitated in 100% TCA/DPBS at −80 °C for 1 hr or overnight. Samples were pelleted at 14,000 rpm for 10 min at 4 °C and resuspended in 500 μL of ice-cold acetone by vortexing. Proteins were pelleted at 5,000 rpm for 10 min at 4 °C. Acetone was removed, and the pellet was left air dry to remove traces of acetone. The pellet was resuspended in 100 μL of DAB buffer (6 M urea, 20 mM Tris-HCl, 1 mM EDTA, 0.5% SDS, pH 8.5) and incubated with 10 mM N-ethylmaleimide (NEM) for 2 hrs at 37 °C with agitation. The reaction was first diluted 3-fold with H_2_O and then 5-fold with ice-cold acetone. Proteins were precipitated at −20 °C for 2 hrs or overnight and pelleted at 4,500 g for 30 min at 4 °C. The protein pellet was re-suspended in 500 μL of ice-cold acetone and centrifuged at 4,500 g for 10 min. Proteins were re-solubilized using 80 μL DAB buffer and incubated with 2.5 mM TCEP for 5 min at 37 °C. The reaction was further diluted with 120 μL DAB buffer and incubated with 10 mM D5-NEM (Cambridge Isotope Laboratories) for 2 hrs at 37 °C. After the final alkylation, samples were diluted 3-fold with H_2_O and then 5-fold with ice-cold acetone for protein precipitation at −20 °C for 2 hrs or overnight. Proteins were pelleted at 4,500 g for 30 min at 4 °C and washed with 500 μL ice-cold acetone. Protein pellets were re-suspended in 200 μL 2 M urea with 1 mM CaCl_2_ and 2 μg trypsin for overnight digestion at 37 °C. 10 μL formic acid was added to the digested peptides. Samples were de-salted using Sep-Pak C18 columns and eluted with 1.5 mL of 80% LC/MS-grade acetonitrile, 20% H_2_O, and 0.1% formic acid. The de-salted samples were dried on SpeedVac for further storage at −20 °C.

Trypsin digested peptide samples were resuspended in 100 μL buffer A peptide concentration was determined using Pierce^TM^ quantitative colorimetric peptide assay. Peptide analysis was performed using an Orbitrap Exploris 240 mass spectrometer coupled to a Dionex Ultimate 3000 RSLCnano system. 500 ng peptides were loaded onto an Acclaim PepMap 100 C18 loading column. Peptides were eluted onto an Acclaim PepMap RSLC column and separated with a 160-min gradient ranging of 5-25% buffer B in buffer A at a flow rate of 0.3 μL/min. The spray voltage was set to 2.1 kV. One full MS scan (350-1,800 m/z, 120,000 resolution, RF lens 65%, automatic gain control (AGC) target 300%, automatic maximum injection mtime, profile mode) was obtained every 2 s with dynamic exclusion (repeat count 2, duration 10 s), isotopic exclusion (assigned), and apex detection (30% desired apex window) enabled. A variable number of MS2 scans (15,000 resolution, AGC 75%, maximum injection time 100 ms, centroid mode was obtained between each MS1 scan based on the highest precursor masses, filtered for monoisotopic peak determination, theoretical precursor isotopic envelope fit, intensity (5E4), and charge state (2-6). MS2 analysis consisted of the isolation of precursor ions (isolation window 2 m/z) followed by higher energy collision dissociation (HCD) (collision energy 30%).

Analysis of MS data collected on Orbitrap Exploris 240 was performed using Thermo Proteome Discoverer software. Protein identification was achieved using the SequestHT and Percolator algorithms^87^ against *Mus Musculus* proteome UniprotKB database.^88^ The protease enzyme was set as trypsin with a maximum of 2 missed cleavages allowed. The peptide precursor mass tolerance was set to 10 ppm with a fragment mass tolerance of 0.02 Da. Dynamic modifications included methionine oxidation (+15.995), N-terminal acetylation (+42.011), methionine loss (−131.040), and cysteine alkylation by either light-NEM (+125.048) or heavy-NEM (+130.079). The FDR for highly confident peptide identification was set at 1%. The resulting H:L ratios were converted to percent oxidation values by the following equation: 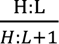. Average percent oxidation values were calculated for each peptide and peptides with standard deviation >50% were removed from the analyses.

### Analysis of OxICAT redox proteomics data

To identify cysteines that were significantly altered in the oxidative proteomics screen, we calculated two metrics, first, change in relative ranks and two, a change in deviation of oxidation. For the first metric, ranks of all the oxidized cysteines were calculated for both WT and NOX1^KO^ and the absolute value of the difference between the two was determined. A change of rank of at least 40 was used to determine the hits. For the second metric, the measure of the deviation from a linear fit with slope 1 and intercept (0,0) was determined. The magnitude of the deviation gives the magnitude of the effect while the direction determines whether it is less or more oxidized in the NOX1^KO^. These two parameters were used for the Volcano plot.

Protein networks that were significantly altered in the different analyses were determined using String-DB.^89^ For determination of up and downregulated canonical pathways, Ingenuity Platform Analysis was used. A cutoff of −log(P-value) >=2 and a Z-score of +/-2 was used for identifying the pathways. A positive z-score implies activation while negative z-score implies inhibition of the pathway. Manually curated subset of pathways and network nodes are shown in the figures.

### IDH1 enzyme activity assay

After recovered from Matrigel, colonoids were washed once with ice-cold PBS and homogenized in 200 μl per well of ice-cold IDH Assay Buffer. Then we centrifuged the lysates at 13000 g for 15 min at 4 °C to get supernatant. The concentrations of supernatant were uniformed to 500 μg/ml by BCA analysis. Then we analyzed IDH enzyme activity following the manufacturer’s instructions.

### αKG levels measurement

After recovered from Matrigel, colonoids were washed once with ice-cold PBS and resuspended in 100 μl per well of ice-cold Assay Buffer. Colon tissues were homogenized in 200 μl of ice-cold Assay Buffer. After incubated on ice for 20 min, we centrifuged the lysates at 13000 g for 15 min at 4 °C to get supernatant. The concentrations of supernatant were uniformed to 1000-2000 μg/ml by BCA analysis. Then we analyzed αKG levels following the manufacturer’s instructions.

### GSH/GSSG ratio detection assay

After recovered from Matrigel, colonoids were washed once with ice-cold PBS and homogenized in 100 μl per well of ice-cold PBS/0.5% NP-40. Then we centrifuged the lysates at 13000 g for 15 min at 4°C to get supernatant. The concentrations of supernatant were uniformed to 500 μg/ml by BCA analysis. After deproteinization, we then ran GSH and Total Glutathione assay following the manufacturer’s instructions.

### Gel electrophoresis and immunoblot analysis

After recovered from Matrigel, colonoids were washed once with cold 1X PBS and were lysed in chilled 1X RIPA buffer (10X stock, EMD Millipore) diluted in PBS containing 1 tablet of cOmplete EDTA free protease inhibitor for 30 minutes on ice. Protein was quantified using a Pierce BCA protein quantification kit. 20 μg of total protein lysates was loaded and separated on Novex™ Tris-Glycine Mini Protein Gels, (4–20%, 1.0 mm, WedgeWell™ format, Thermo Scientific). Proteins were transferred to nitrocellulose membranes, blocked with EveryBlot blocking buffer (Bio-Rad), and incubated with the indicated primary antibodies diluted in blocking buffer at 4 °C overnight. Membranes were washed three times in TBS-T and incubated with HRP-coupled secondary antibodies for 1 h at room temperature. After washed for three times, proteins on the membranes were detected by chemiluminescence using ECL (Thermo Scientific) in a Bio-Rad ChemiDoc Imager. The following primary antibodies and dilutions were used: anti-HIF1α (NOVUS Biological, NB100-134SS, 1:1000), anti-HIF1α (CST, 36169S, 1:1000) and anti-β-actin (A5441, Sigma, 1:1000). Western blot images were processed using FIJI and Affinity Designer.

### H&E analysis, AB/PAS analysis and DAB analysis of HIF1α

Mice were euthanized and colon tissues were collected and fixed with 4% paraformaldehyde diluted in PBS at 4 °C overnight. Tissues were processed, paraffin-embedded, and cut into 5 μm sections and under H&E (hematoxylin and eosin), AB/PAS (Alcian blue/Periodic acid-Schiff) staining at BIDMC histology core. Colon FFPE samples were deparaffinized, rehydrated and performed with antigen retrieval and we then followed the manufacturer’s instructions for the DAB analysis (VECTASTAIN® Elite® ABC-HRP Kit and ImmPACT® DAB Substrate Kit). The following primary antibodies and dilutions were used: anti-HIF1α (NOVUS Biological, NB100-134SS, 1:1000). Whole cross-sections were tile-scanned and imaged on a widefield BioTek Cytation5 automated microscope at 20X, and images were analyzed with FIJI.

### ChIP-qPCR

ChIP-qPCR was performed as described previously with some modifications from Shen et al.^90^ Briefly, 2D monolayer cells of human colonoids H514 were crosslinked with 1% formaldehyde for 10 min at room temperature and quenched by 2.5 M glycine for 5 min. Cells were lysed with nuclei lysis buffer and lysates were sonicated to an average chromatin fragment of 300-400 bp. Take 10% out as input. The following antibodies were used for each ChIP experiment based on instructions: 1:50 dilution anti-HIF1α (Novus Biologicals, NB100-134), 1:50 dilution anti-TCF4 (CST, #2569T). 40 μl of Protein G dynabeads were used for each ChIP experiment. The purified elution was used for following qPCR with primers listed in Key sources table.

### Quantification and data analysis

For all quantifications, the exact value of n is described in detail in the Figure legends, where n indicates biological replicates, either number of mice, number of wells containing cells or number of areas containing cells. For all microscopy analysis, images were blinded prior to scoring and quantification. All statistical analyses were performed using Microsoft Excel and GraphPad Prism software. Data were represented as mean ± SEM throughout the Figures unless indicated and the level of significance was indicated by asterisks for the following corresponding p-values: _∗_p < 0.05, ^∗∗^p < 0.01, ^∗∗∗^p < 0.001, ^∗∗∗∗^p < 0.0001. The specific statistical test used to for each experiment is described in detail in the Figure legends. For comparisons between two groups, we performed student’s two-sided t-tests. For comparisons that have more than two groups or two conditions, One-way ANOVA and Two-way ANOVA were utilized. Sample sizes for all experiments were chosen according to standard practice in the field.

